# Central and peripheral GLP-1 systems independently and additively suppress eating

**DOI:** 10.1101/2020.08.03.234427

**Authors:** Daniel I. Brierley, Marie K. Holt, Arashdeep Singh, Alan de Araujo, Macarena Vergara, Majd H. Afaghani, Shin Jae Lee, Karen Scott, Wolfgang Langhans, Eric Krause, Annette de Kloet, Fiona M. Gribble, Frank Reimann, Linda Rinaman, Guillaume de Lartigue, Stefan Trapp

## Abstract

The anorexigenic peptide glucagon-like peptide-1 (GLP-1) is secreted from gut enteroendocrine cells and brain preproglucagon (PPG) neurons, which respectively define the peripheral and central GLP-1 systems. As peripheral satiation signals are integrated in the nucleus tractus solitarius (NTS), PPG^NTS^ neurons are assumed to link the peripheral and central GLP-1 systems, forming a unified GLP-1 gut-brain satiation circuit. This hypothesis, however, remains unsubstantiated. We report that PPG^NTS^ neurons encode satiation in mice, consistent with vagal gastrointestinal distension signalling. However, PPG^NTS^ neurons predominantly receive vagal input from oxytocin receptor-expressing vagal neurons, rather than those expressing GLP-1 receptors. Furthermore, PPG^NTS^ neurons are not necessary for eating suppression induced by the GLP-1 receptor agonists liraglutide or semaglutide, and semaglutide and PPG^NTS^ neuron activation additively suppress eating. Central and peripheral GLP-1 systems thus suppress eating via independent gut-brain circuits, hence PPG^NTS^ neurons represent a rational pharmacological target for anti-obesity combination therapy with GLP-1 receptor agonists.

Graphical Abstract:

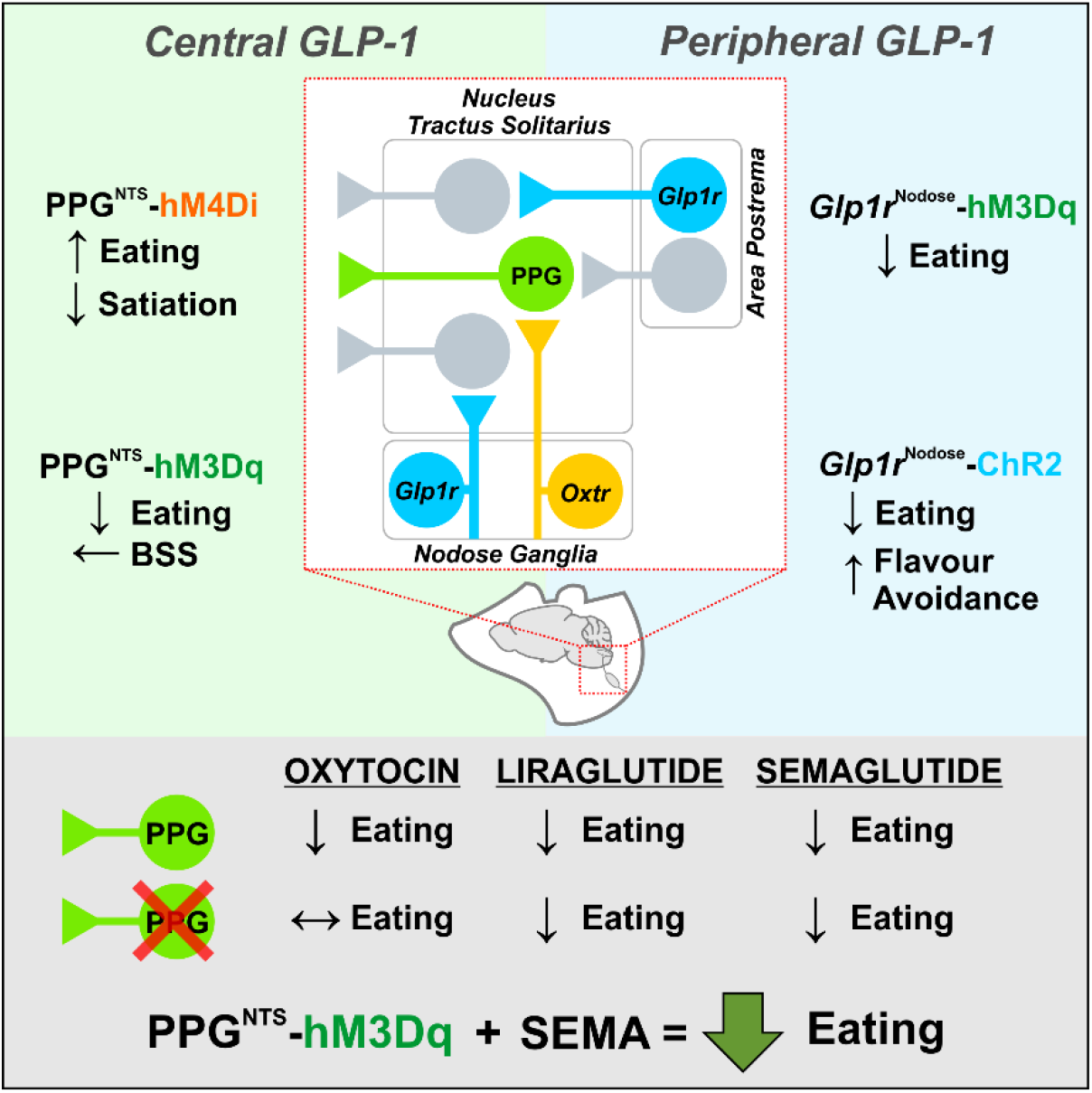

---

Glucagon-like peptide 1 (GLP-1) acts as an incretin hormone and anorexigenic neuropeptide, prompting the successful and ongoing development of GLP-1-based therapies for type 2 diabetes and obesity^1,2^. Endogenous GLP-1 is produced both by enteroendocrine cells in the gut, and preproglucagon (PPG) neurons in the brain, which are the defining populations of the peripheral and central GLP-1 systems, respectively^3,4^. PPG neurons in the nucleus tractus solitarius (PPG^NTS^ neurons) suppress eating when chemogenetically or optogenetically activated^5–7^, consistent with substantial pharmacological evidence for anorexigenic GLP-1 signalling in the brain (reviewed by Muller et al^4^). Physiologically, PPG^NTS^ neurons are the major source of GLP-1 in the brain, are necessary for stress-induced hypophagia, and their inhibition or ablation elicits transient hyperphagia during large intakes^7^. Although glutamate is a co-transmitter in these neurons^8,9^, selective *Ppg* knockdown has confirmed the necessity of proglucagon-derived peptides for their anorexigenic role^10^. PPG^NTS^ neurons are thus the crucial component of the central GLP-1 system, which they comprise along with numerous populations of GLP-1 receptor (GLP-1R)-expressing neurons found throughout the brain^11,12^. Whilst it is widely assumed that endogenous *peripheral* GLP-1 interacts with the *central* GLP-1 system (via vagal and/or endocrine activation of PPG^NTS^ neurons) to control eating under physiological conditions, this link remains subject to debate and has not been demonstrated empirically^4,13–15^. The difficulty in substantiating this link partly arises from the inherent complexity of interrogating these widely-distributed systems using pharmacological approaches, compounded by observations that native GLP-1 and pharmacokinetically-optimized therapeutic GLP-1 receptor agonists (GLP-1RAs) suppress eating via divergent signalling pathways^16–22^.

Numerous studies demonstrate the value of selective transgenic manipulations to determine the neuroanatomical organization and physiological functions of specific cell populations involved in eating control, including those comprising parts of the peripheral and central GLP-1 systems^7,23–26^. Here we utilized similar transgenic and viral approaches to address whether PPG^NTS^ neurons have a role in physiological satiation, and determined their anatomical and functional connectivity to molecularly defined neuronal populations mediating gut-brain satiation signalling. Specifically, we tested the prevalent but unsubstantiated hypothesis that peripheral GLP-1 signals to the brain to suppress eating via vagal and/or endocrine activation of central GLP-1-producing preproglucagon (PPG) neurons, i.e. that peripheral and central GLP-1 systems comprise a unified, directly connected gut-brain satiation circuit. We furthermore tested the role of PPG^NTS^ neurons in eating suppression induced by the anti-obesity GLP-1RAs liraglutide and semaglutide, to establish whether this neuronal population has translational importance as a distinct therapeutic target for obesity treatment.

## Results

### PPG^NTS^ neurons selectively encode large meal satiation

PPG^NTS^ neurons are not necessary for control of daily or long term food intake or bodyweight in *ad libitum* eating mice^7^. However, it is unknown whether they regulate within- or between-meal parameters, or whether the absence of an ablation-induced bodyweight phenotype masks more subtle alterations in energy expenditure or physical activity. We addressed these questions by metabolic phenotyping of *ad libitum* eating mice following viral ablation of PPG^NTS^ neurons (Fig. S1a). Food intake over the circadian cycle was unaffected by neuronal ablation in either sex (Fig. S1b-c,k-l), and there was no effect on meal size, frequency or duration (Fig. S1e-g). Similarly, ablation did not affect locomotion, energy expenditure, bodyweight or water intake (Fig. S1d, i,j,m). As a positive control in this model we did, however, successfully replicate a previous report^7^ that ablation induces hyperphagia both during post-fast refeeding, and after a short liquid diet preload (Fig. S1n-o).

The replicable observation that ablation of PPG^NTS^ neurons elicits transient hyperphagia only under conditions manipulated to induce large intakes^7^ is consistent with results in rats indicating that these neurons are activated during ingestion of unusually large meals^27^. We thus tested the hypothesis that hyperphagic responses observed after PPG^NTS^ neuronal ablation are specifically due to a delay in meal termination, to establish a *bona fide* role for PPG^NTS^ neurons in the process of satiation. We conducted high resolution meal pattern analysis using home cage FED pellet dispensers (Feeding Experimentation Device^28^), and observational analysis of liquid diet intake, to test the effects of acute chemogenetic inhibition of PPG^NTS^ neurons on termination of large solid and liquid meals (Fig. 1a,g). PPG^NTS^ inhibition increased fasting-induced pellet refeeding during hour 1 in a sex-independent manner, and this effect was driven by increased meal size, rather than frequency (Fig. 1b-f). The hyperphagic effect in this model was also confirmed to be specific to large meals, as inhibition had no effect under *ad libitum* eating conditions (Fig. S1p-r). We then modified a previously used Ensure liquid diet preload paradigm^7^ to test whether PPG^NTS^ neurons are necessary for satiation during consumption of large liquid meals (Fig. 1g), as suggested by a previous cFos expression study in rats^29^. Under these conditions, Ensure intake was increased by PPG^NTS^ neuron inhibition, and, consistent with effects on pellet intake in the FED system, this increase was driven by increased duration of Ensure eating (Fig. 1h-k). These data demonstrate that both under conditions of normal and negative energy balance, PPG^NTS^ neurons are recruited to encode physiological satiation specifically by ingestion of large meals.

**Figure 1.**
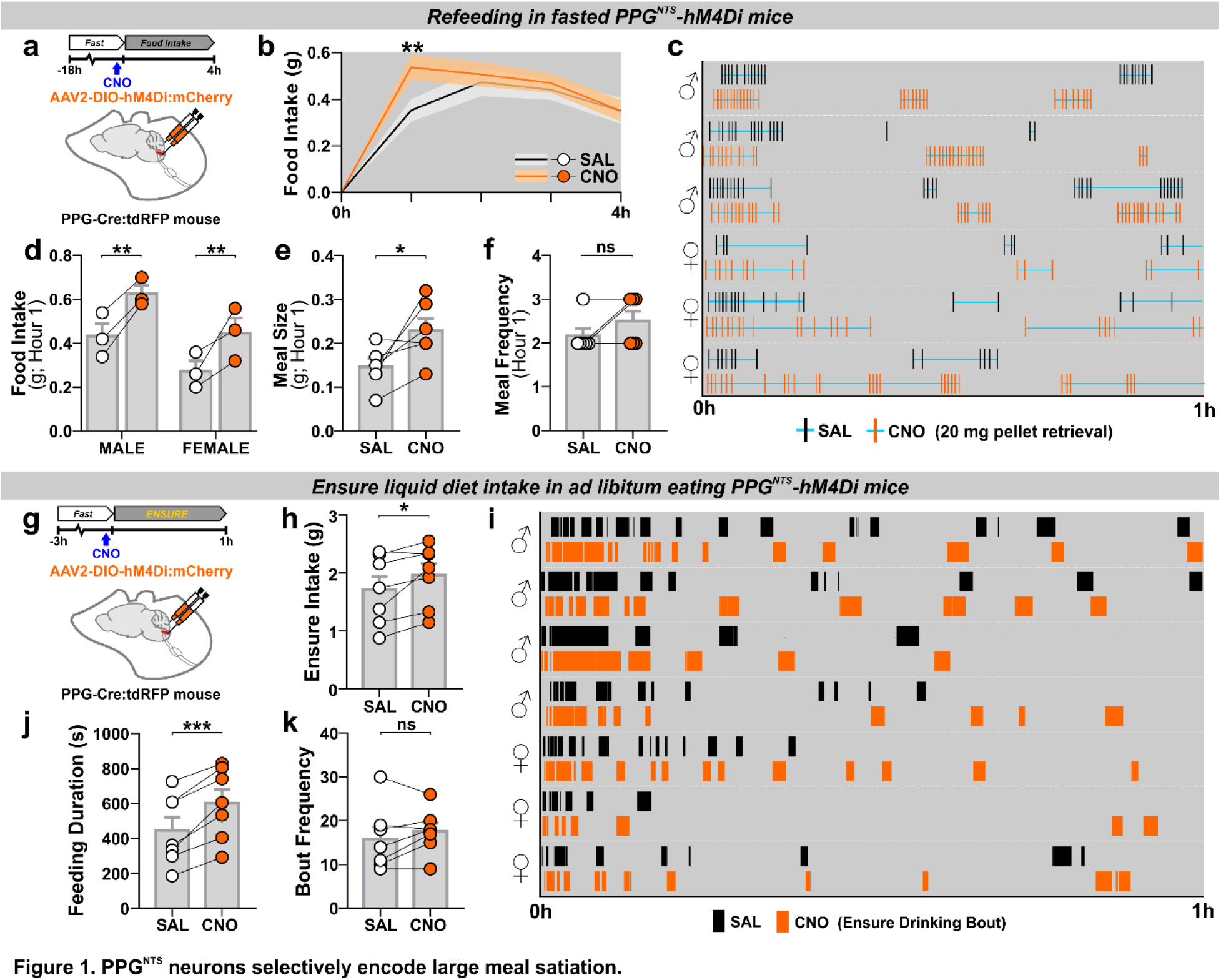
PPG^NTS^ neurons selectively encode large meal satiation. (a) Experimental model and paradigm for meal pattern analysis of post-fast refeeding in PPG^NTS^-hM4Di mice (n=6) using FED system. (b) 4h dark phase food intake, 2-way within-subjects ANOVA: Drug × Time *F*_(3,15)_=3.664, *p*=0.0367. (c) Raster plot of chow pellet retrievals over 1h dark phase. Plots from the same mouse after saline and CNO injections presented adjacently. (d) 1h intake by sex, 2-way mixed-model ANOVA: Drug *F*_(1,4)_=29.09, *p*=0.0057. (e-f) Meal pattern parameters during 1h refeed, paired 2-tailed t-test or Wilcoxon matched-pairs test: E) *t*_(5)_=2.757, *p*=0.040; F) *W*=−3, *p*=0.500. (g) Experimental model and paradigm for temporal analysis of Ensure intake in PPG^NTS^-hM4Di mice (n=7). (h) 1h Ensure intake, paired 2-tailed t-test: *t*_(6)_=2.859, *p*=0.0288. Ensure intake was sex-independent (data not shown), 2-way mixed-model ANOVA: Drug *F*_(1,5)_=11.58, *p*=0.0192. (i) Raster plot of Ensure drinking bouts over 1h dark phase. Plots from the same mouse after saline and CNO injections presented adjacently. (j-k) Temporal parameters of Ensure drinking, paired 2-tailed t-test: J) *t*_(6)_=6.55, *p*=0.0006; K) *t*_(6)_=1.263, *p*=0.254; M) *t*_(6)_=4.784, *p*=0.0031.

### PPG^NTS^ neurons suppress eating without behavioural disruption

The observation that PPG^NTS^ neurons selectively encode satiation during large meals, but apparently do not control intake under *ad libitum* eating conditions, suggests they have capacity to suppress eating when stimulated. Evidence for such capacity has been reported previously^5–7^, and could indicate translational potential for PPG^NTS^ neurons as a target for therapeutic suppression of eating, provided that the eating suppression is robust, is not compensated for, and is not associated with nausea/malaise. We therefore extended these studies by testing whether hypophagia induced by chemogenetic activation of PPG^NTS^ neurons was followed by compensatory rebound hyperphagia, or elicited significant disruption to the behavioural satiety sequence (Fig. 2a,e). In *ad libitum* eating mice, PPG^NTS^ activation reduced intake by ~40% in the first 24 hours (Fig. 2b) in a sex-independent manner (Fig. S2b), predominantly driven by reductions in the first 5 hours of the dark phase (Fig. S2a). No compensatory hyperphagia occurred, hence cumulative intake and bodyweight were still reduced 48 hours after acute CNO administration (Fig. 2c-d). This robust suppression of eating with sustained reduction in intake was also observed when PPG^NTS^ neurons were activated immediately prior to dark onset refeeding after a prolonged (18hr) fast (Fig. 2e, S2f). We thus combined the FED system with infrared video in this paradigm to investigate changes to the behavioural satiety sequence under relatively naturalistic home cage conditions. Following a period of eating (Fig. S2g), PPG^NTS^ neuron activation advanced the point of satiation by ~15 minutes (shift from time bin 4 to 1; Fig. 2f-g), and the stochastic sequence of satiety behaviours (eating → grooming → inactive) was not disrupted. Quantitative analyses revealed that PPG^NTS^ activation did not significantly alter eating rate (Fig. S2h), but reduced eating duration during the first 15 minutes (Fig. 2h), as expected from the left-shifted satiation point. Time inactive was modestly increased, and grooming and active behaviours appeared to be correspondingly decreased, however the temporal patterns of these behaviours were maintained (Fig. 2i-k). That the robust suppression of intake elicited by their activation is not associated with significant alteration to the behavioural satiety sequence further supports the idea that PPG^NTS^ neurons have translational potential as a pharmacological target for eating suppression. These observations also contrast with previously reported effects in this assay of the emesis/nausea-inducing agent lithium chloride and the GLP-1RA Exendin-4, which reduce eating rate and almost completely suppress grooming and other active behaviours^30,31^. Instead, and consistent with prior evidence that PPG^NTS^ activation does not condition flavour avoidance^5^, the absence of behavioural satiety sequence disruption reported here supports the view that PPG^NTS^ neuronal activation suppresses eating without inducing nausea/malaise, in contrast to the effects of peripherally administered GLP-1RAs.

**Figure 2.**
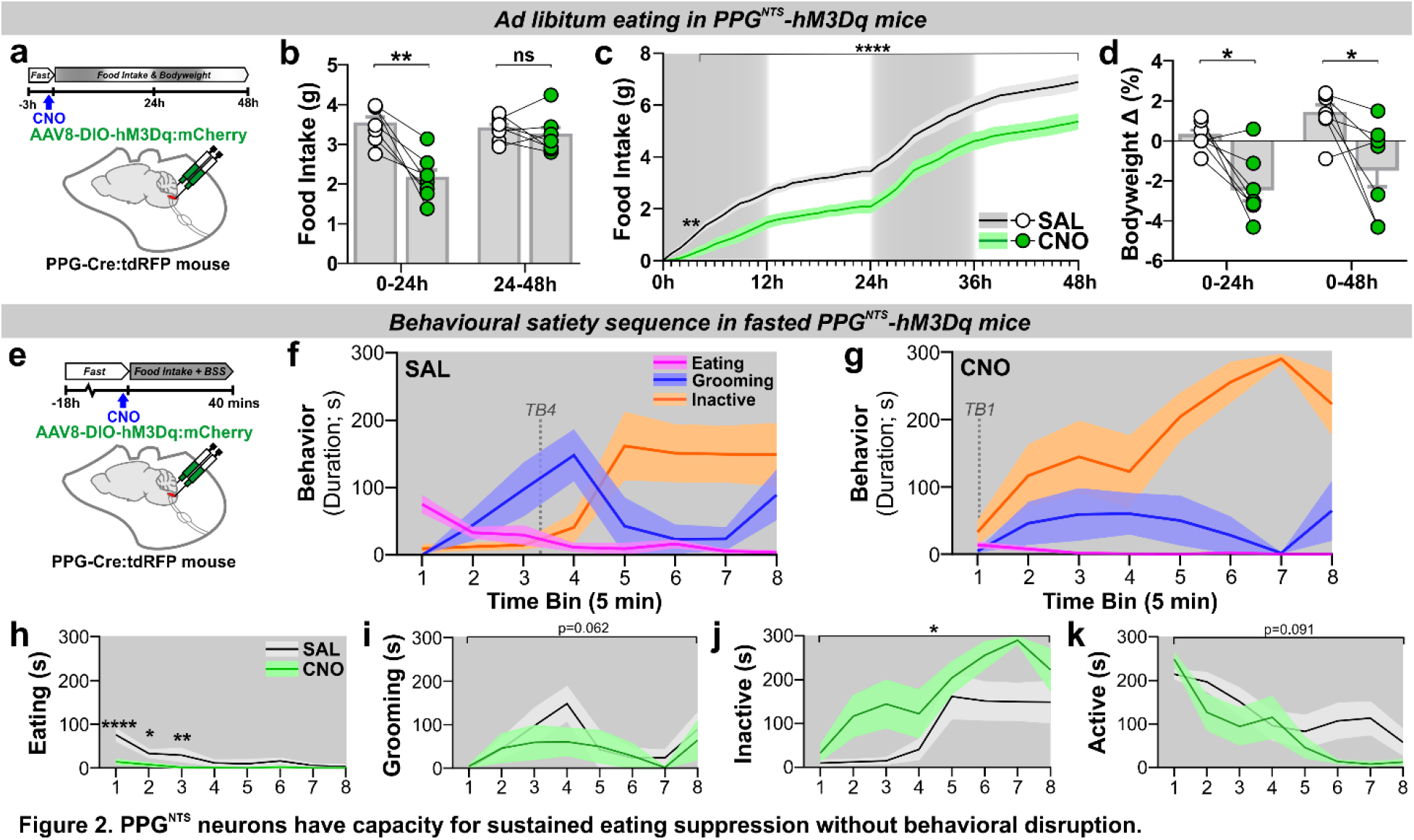
PPG^NTS^ neurons have capacity for sustained eating suppression without behavioural disruption. (a) Experimental model and paradigm for *ad libitum* pellet eating from FED in PPG^NTS^-hM3Dq mice (n=7). (b) Daily food intake during 48h test, 2-way within-subjects ANOVA: Drug × Day *F*_(1,6)_=14.52, *p*=0.0089. (c) Cumulative hourly food intake over two days, 2-way within-subjects ANOVA: Drug × Time *F*_(48,288)_=6.481, *p*<0.0001. (d) 24h and 48h bodyweight change, 2-way within-subjects ANOVA: Drug *F*_(1,6)_=10.41, *p*=0.018. (e) Experimental model and paradigm for BSS analysis in 18h fasted PPG^NTS^-hM3Dq mice (n=7). (f-g) Behavioural satiety sequences following saline and CNO injections. Satiation point/satiety onset (when duration inactive exceeds eating) shown by dotted lines. (h-k) Quantitative analysis of hM3Dq effect on BSS behaviours, 2-way with-subjects ANOVA: h) Drug x Time *F*_(7,42)_=5.673, *p*=0.0001; i) Drug *F*_(1,6)_=5.261, *p*=0.0616; j) Drug *F*_(1,6)_=12.48, *p*=0.0123; k) Drug *F*_(1,6)_=4.028, *p*=0.0915.

### *Glp1r*-expressing vagal afferent neurons suppress eating and condition flavour avoidance

Direct synaptic input from undefined population(s) of vagal afferent neurons (VANs) to PPG^NTS^ neurons^26,32^ presumably underlies the ability of gastrointestinal distension and large liquid intakes to induce cFos expression in this NTS population^29,33^, and to drive their role in large meal satiation. However, the molecular identities of VAN inputs to PPG^NTS^ neurons remain to be characterized. VANs defined by their expression of the GLP-1 receptor gene (*Glp1r*) innervate the gut and have been identified as a predominantly mechanosensory population that encodes gastrointestinal distension, as well as likely mediating paracrine satiation signalling by peripheral GLP-1^14,24,25^. However, it is unknown whether PPG^NTS^ neurons are a major synaptic target of *Glp1r* VANs, and thus to what extent direct vagal communication between peripheral and central GLP-1 systems is neuroanatomically plausible. To address this question, we developed approaches for viral targeting and activation of *Glp1r* VANs with chemogenetic and optogenetic effectors (Fig. 3a,f), to determine whether these manipulations produced effects on eating consistent with *Glp1r* VANs being part of a unified gut-brain satiation circuit with PPG^NTS^ neurons. Chemogenetic activation of *Glp1r* VANs in *ad libitum* eating mice suppressed eating during the dark but not subsequent light phase (Fig. 3b, S3a). Notably, however, the magnitude of this hypophagic effect appeared less robust than that following chemogenetic activation of PPG^NTS^ neurons. Bodyweight was also transiently decreased, driven by suppressed eating rather than increased energy expenditure (Fig. 3c-e, S3b-c). The modest anorexigenic effect of chemogenetically activating *Glp1r* VANs in *ad libitum* eating mice precluded use of the 18hr fasted BSS paradigm in this model. We therefore instead tested whether optogenetic activation of the central axon terminals of ChR2-transduced *Glp1r* VANs was able to condition avoidance of, or a preference for, a paired novel flavour of Kool-Aid (Fig. 3f-g). Optogenetic activation of *Glp1r* terminals within the NTS conditioned avoidance of the paired flavour (Fig. 3h), and also modestly suppressed eating in a subsequently conducted acute eating test (Fig. 3i). This finding contrasts with the lack of disruption to the BSS we observed following activation of PPG^NTS^ neurons, and a previous report that chemogenetic activation of PPG^NTS^ neurons does not condition flavour avoidance^5^, but is consistent with several reports that GLP-1RAs condition flavour avoidance and reduce reward-related behaviours^34–36^. These results support prior evidence that *Glp1r* VANs at least partly mediate endogenous or exogenous peripheral GLP-1 satiation signalling^20,22^. However, the modest anorexigenic effect and conditioning of avoidance produced by chemogenetic and optogenetic activation of *Glp1r* VANs argues against the hypothesis that this population are the primary driver of eating suppression by PPG^NTS^ neurons.

**Figure 3.**
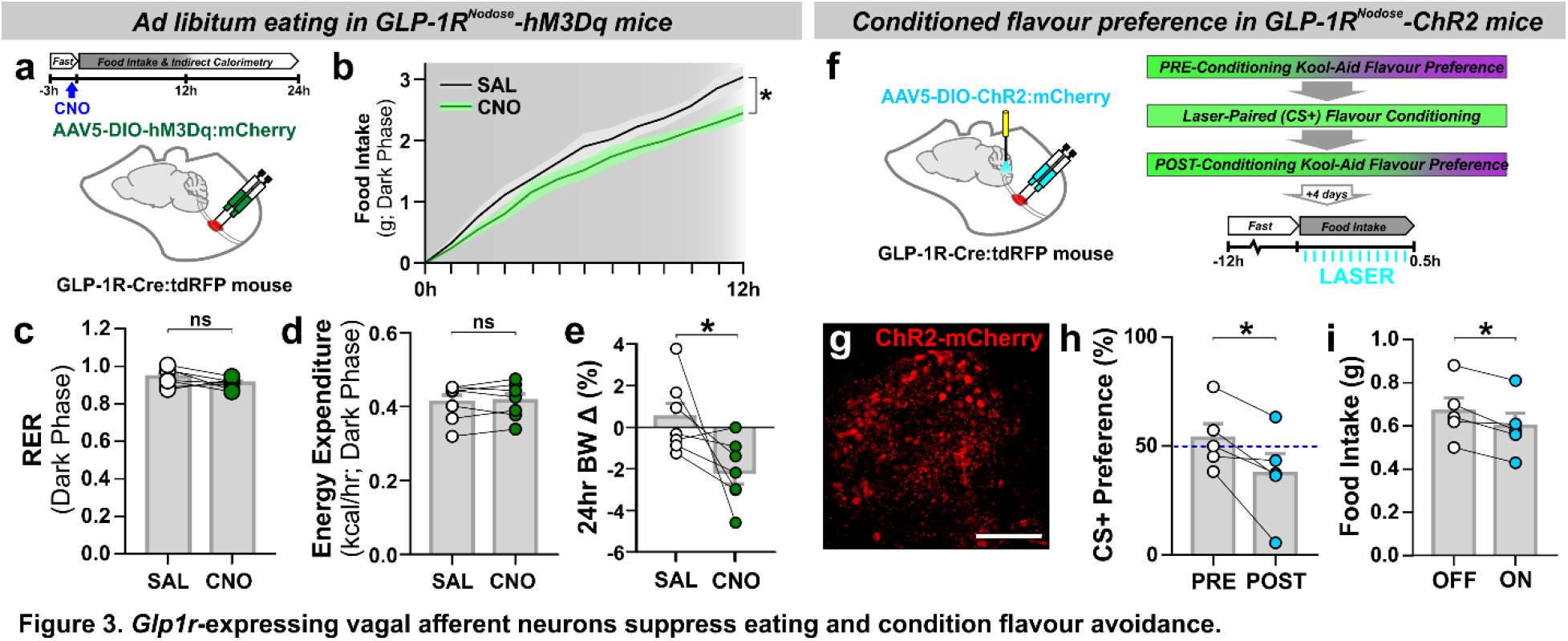
*Glp1r*-expressing vagal afferent neurons suppress eating and condition flavour avoidance. (a) Experimental model and paradigm for food intake and metabolic analysis of *ad libitum* eating GLP-1R^Nodose^-hM3Dq mice (n=7). (b) Cumulative hourly dark phase food intake, 2-way within-subjects ANOVA: Drug × Time *F*_(12,144)_=2.078, *p*=0.0218. (c-e) Dark phase metabolic parameters and 24h bodyweight change, paired 2-tailed t-test: c) *t*_(6)_=1.642, *p*=0.152; d) *t*_(6)_=0.543, *p*=0.607; e) *t*_(6)_=2.323, *p*=0.0296. (f) Experimental model and paradigm for optogenetically-evoked conditioned flavour preference and intake analysis in GLP-1R^Nodose^-ChR2 mice (n=5). (g) Z-projection photomicrograph of ChR2-mCherry expression in nodose ganglia. Scale=100μm. (h-i) Conditioned stimulus (CS+) preference and 0.5h food intake, paired 2-tailed t-test: h) *t*_(4)_=3.216, *p*=0.0324; i) *t*_(4)_=3.976, *p*=0.0165.

### *Oxtr* rather than *Glp1r* VANs are the major vagal input to PPG^NTS^ neurons

We next tested the neuroanatomical connectivity between PPG^NTS^ neurons and *Glp1r* VANs, using two complementary circuit mapping approaches. Utilizing a cross of GLP-1R-Cre and PPG-YFP mouse strains^37^ combined with unilateral viral targeting of nodose ganglia, we selectively labelled the NTS terminal fields of *Glp1r* VANs with tdTomato, allowing simultaneous visualization with YFP-expressing PPG^NTS^ somata and dendrites (Fig. 4a). While extensive innervation by both right and left branch *Glp1r* VANs was observed along the rostro-caudal extent of the NTS, there was little regional overlap with PPG^NTS^ neurons. In the caudal NTS, *Glp1r* vagal afferents predominantly terminated dorsomedial to PPG^NTS^ somata, and their terminal fields extended considerably beyond the rostral extent of the PPG^NTS^ population (Fig. 4b-c).

**Figure 4.**
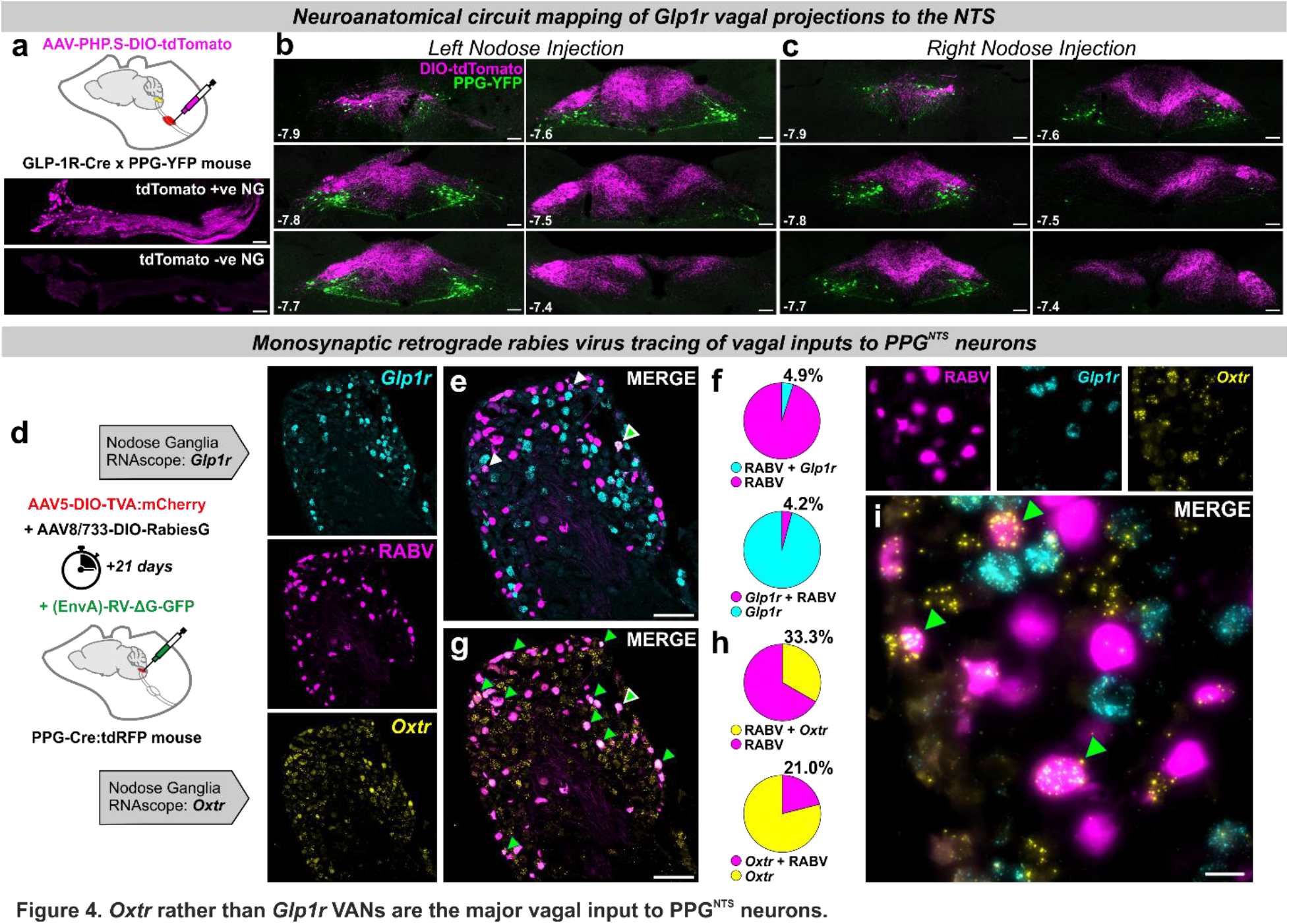
*Oxtr* rather than *Glp1r* VANs are the major vagal input to PPG^NTS^ neurons. (a) Experimental model for viral-mediated mapping of left and right *Glp1r* vagal afferent projections to the NTS, and representative photomicrographs of tdTomato expression in virus-injected nodose ganglia (NG) and non-injected contralateral NG (n=3 mice/side). Scale=100μm. (b-c) Photomicrographs of tdTomato-expressing terminal fields of L and R branch *Glp1r* vagal afferents along the rostro-caudal extent of the NTS (mm posterior to Bregma in bottom left) in PPG-YFP mice. Scale=100μm. (d) Experimental model for rabies virus (RABV)-mediated monosynaptic retrograde tracing of vagal inputs to PPG^NTS^ neurons combined with RNAscope fluorescence *in situ* hybridization (FISH) for GLP-1R (*Glp1r*) and oxytocin receptor (*Oxtr*) transcripts (n=8 mice). (e) Photomicrograph of nodose ganglion showing rabies virus GFP expression (RABV) and *Glp1r* FISH. RABV+*Glp1r* co-localization shown by white arrows, RABV+*Glp1r*+*Oxtr* by white-edged green arrow. Scale=100μm. (f) RABV and *Glp1r* co-localization as proportions of all RABV+ cells and all *Glp1r*+ cells, from 903 RABV+, 1188 *Glp1r*+ and 1460 *Oxtr*+ cells from L and R NG. (g) Photomicrograph of nodose ganglion showing RABV expression and *Oxtr* FISH. Scale=100μm. (h) RABV and *Oxtr* co-localization as proportions of all RABV+ cells and all *Oxtr*+ cells. RABV+*Oxtr* co-localization shown by green arrows, RABV+*Glp1r*+*Oxtr* by white-edged green arrow. (i) High magnification Z-projection of RABV, *Glp1r* and *Oxtr* cells in NG. Scale=20μm.

The absence of overlap between *Glp1r* VAN terminals and PPG^NTS^ somata does not preclude some vagal input via their distal dendrites in the dorsomedial NTS. We therefore quantified direct synaptic connectivity between *Glp1r* VANs and PPG^NTS^ neurons by Cre-dependent monosynaptic retrograde rabies virus tracing in combination with RNAscope fluorescence *in situ* hybridization for *Glp1r* and oxytocin receptor gene (*Oxtr*) expression in nodose ganglia (Fig. 4d). Expression of *Oxtr* was investigated based on reports of target-based scRNAseq analysis of VANs (target-scSeq), which suggest that mechanosensation of gastric and intestinal distension are predominantly encoded by VANs defined by *Glp1r* and *Oxtr* expression, respectively (with some overlap), and that an additional subpopulation of chemosensitive VANs expressing *Glp1r* (but not *Oxtr*) innervate the intestine and mediate paracrine GLP-1 satiation signalling^14,25^. As previously reported^26^, we observed that rabies virus-GFP was expressed extensively in nodose ganglia, confirming substantial monosynaptic vagal innervation of PPG^NTS^ neurons. Surprisingly, however, we found that <5% of PPG^NTS^ neuron-innervating VANs express *Glp1r* alone, and similarly <5% of VANs expressing *Glp1r* alone synapse onto PPG^NTS^ neurons (Fig. 4e-f). Conversely, 33% of all PPG^NTS^ neuron-innervating VANs express *Oxtr* alone, and 21% of VANs which express *Oxtr* alone synapse onto PPG^NTS^ neurons (Fig. 4g-h). VANs expressing both *Oxtr* and *Glp1r* (which are presumably mechanosensory) comprise 20-25% of these populations (Fig. S4c), and 26% of these *Oxtr* / *Glp1r* VANs synapse onto PPG^NTS^ neurons, comprising an additional 9% of vagal input to this population (Fig. S4d-h). We thus identified PPG^NTS^ neurons as an important synaptic target of *Oxtr* VANs, in addition to the catecholaminergic target population previously identified^25^. Our findings strongly suggest that *Oxtr*-rather than *Glp1r*-expressing VANs are the primary source of gastrointestinal distension signals driving PPG^NTS^ neuron-mediated satiation. Furthermore, as the overwhelming majority of VANs expressing *Glp1r* but not *Oxtr* (which presumably includes the chemosensory population) do not synapse onto PPG^NTS^ neurons, they are highly unlikely to be a functionally relevant target of vagal-dependent paracrine signalling from the peripheral GLP-1 system.

### PPG^NTS^ neurons are necessary for oxytocin-induced eating suppression

Having identified *Oxtr*-expressing VAN input to PPG^NTS^ neurons, we next characterized the effects of oxytocin itself on this population. We first performed *ex vivo* calcium imaging using coronal brainstem slices from transgenic mice expressing GCaMP3 in PPG^NTS^ neurons^38,39^. Slices were taken at a rostro-caudal level containing the majority of PPG^NTS^ neurons, and which is reported to contain substantial *Oxtr* VAN terminal fields^25^. Superfusion of oxytocin activated 84% of glutamate-responsive PPG^NTS^ neurons, as determined by increased calcium-dependent fluorescence (Fig. 5a-e). Notably, superfusion of GLP-1 has no effect on PPG^NTS^ neurons in the same preparation^37^.

**Figure 5.**
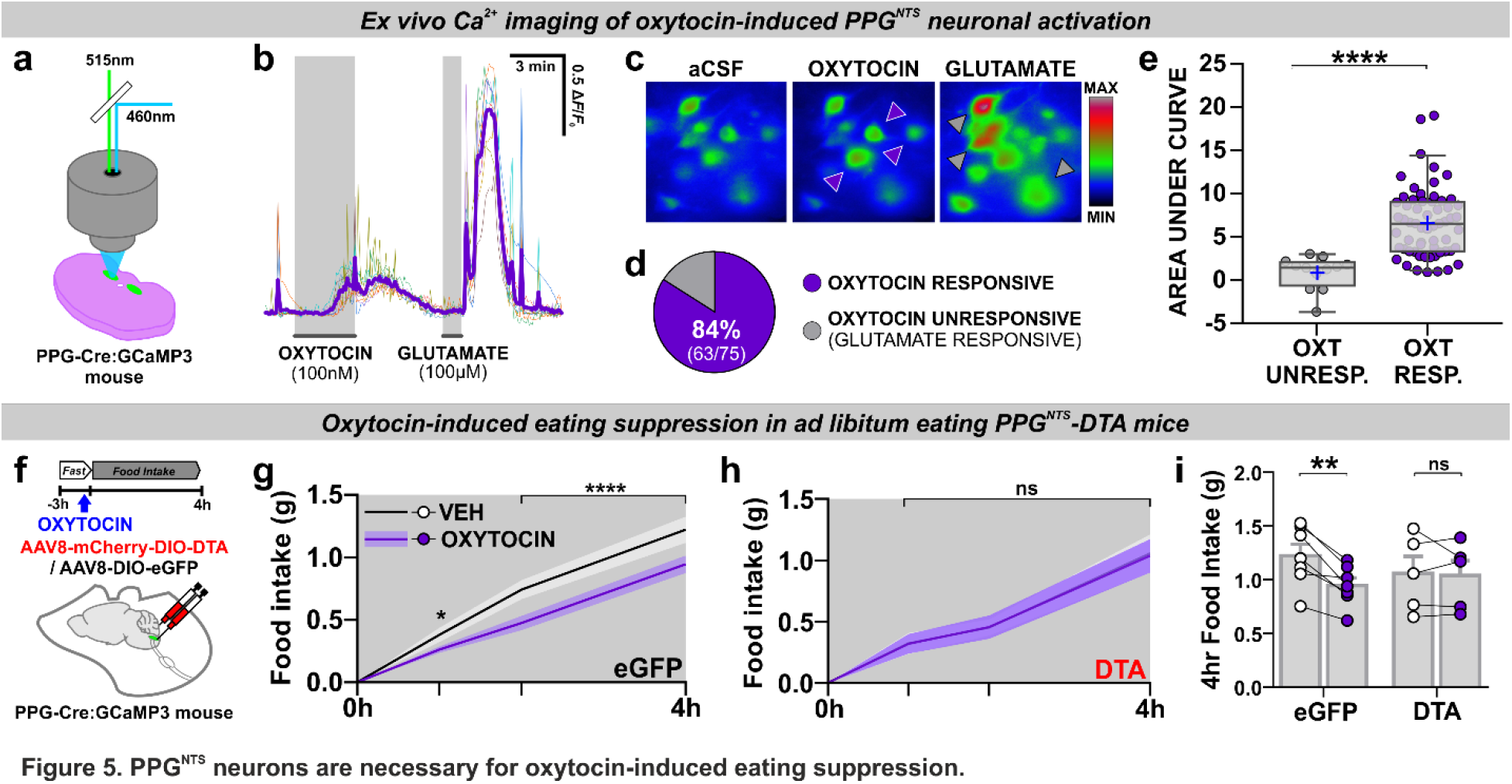
PPG^NTS^ neurons are necessary for oxytocin-induced eating suppression. (a) Experimental model for imaging of oxytocin-induced neuronal calcium dynamics in *ex vivo* brainstem slices from mice expressing GCaMP3 in PPG neurons (n=75 cells from 3 mice). (b) Representative Δ*F*/*F*_0_ traces from individual neurons and mean response (purple line) during bath application of oxytocin and glutamate. (c) Representative images of PPG^NTS^:GCaMP3 neurons pseudocolored for fluorescence intensity under baseline conditions (aCSF) and responding to oxytocin (purple arrows) and glutamate (grey arrows). (d) Oxytocin-responsive PPG^NTS^:GCaMP3 neurons as a proportion of all glutamate-responsive PPG^NTS^ neurons (i.e. healthy neurons with functional GCaMP3 expression). (e) Median AUC during exposure to oxytocin in oxytocin unresponsive and responsive PPG^NTS^:GCaMP3 neurons, Mann-Whitney 2-tailed U-test: U=48, *p*<0.0001. (f) Experimental model and paradigm for oxytocin-induced eating suppression in PPG^NTS^-DTA ablated mice (DTA; n=5) or eGFP-transduced controls (eGFP; n=7). (g-h) Cumulative 4h dark phase food intake in eGFP and DTA mice administered oxytocin (0.4 mg/kg, i.p.), 2-way within-subjects ANOVA: g) Drug × Time *F*_(2,12)_=6.133, *p*=0.0146; h) Drug *F*_(1,4)_=0.0117, *p*=0.919. (i) 4h food intake by virus, 2-way mixed-model ANOVA: Drug × Virus *F*_(1,10)_=8.472, *p*=0.0155.

Peripherally administered oxytocin is reported to suppress eating in a vagal-dependent manner^40,41^, so we subsequently tested whether PPG^NTS^ neurons were necessary for this effect, given their direct synaptic inputs from *Oxtr* VANs. While oxytocin acutely suppressed eating in *ad libitum* eating control mice, this effect was completely abolished in PPG^NTS^ neuron-ablated mice (Fig. 5f-i, S5a-b), confirming these neurons as a necessary component of the gut-brain circuit recruited by peripheral oxytocin to suppress eating.

### PPG^NTS^ neurons are not a major synaptic target of area postrema *Glp1r* neurons

Having determined the neuroanatomical implausibility of vagal transmission of peripheral GLP-1 signals to PPG^NTS^ neurons, we next investigated a potential route for endocrine GLP-1 signalling to these NTS neurons via *Glp1r* neurons in the area postrema (AP), which lacks a blood-brain barrier and has been implicated as a site where circulating GLP-1 and GLP-1RAs may act to suppress eating^17,20,42,43^. Monosynaptic retrograde tracing from PPG^NTS^ neurons combined with *in situ* hybridization for *Glp1r* was again utilized (Fig. 6a). In contrast to robust synaptic input from VANs, synaptic inputs to PPG^NTS^ neurons from the AP were relatively sparse (Fig. 6b). Of these sparse inputs from the AP, 25% expressed *Glp1r*, however these represented <3% of all *Glp1r* AP neurons (Fig. 6c). As catecholaminergic AP neurons express GLP-1R and have been proposed to link peripheral GLP-1 signalling to central nuclei (including the NTS) involved in eating control^43^, we further characterized PPG^NTS^ neuronal input from tyrosine hydroxylase immunoreactive AP neurons (Fig. 6d-e, S6a-c). ~20% of the sparse AP inputs to PPG^NTS^ neurons are catecholaminergic, consistent with a report that PPG^NTS^ neurons are indirectly activated by noradrenaline^44^. However, these presynaptic AP neurons comprised only 3% of all catecholaminergic AP neurons (Fig. 6e). Therefore, PPG^NTS^ neurons are unlikely to be a functionally relevant target for peripheral GLP-1 and/or GLP-1RAs acting via the AP to suppress eating.

**Figure 6.**
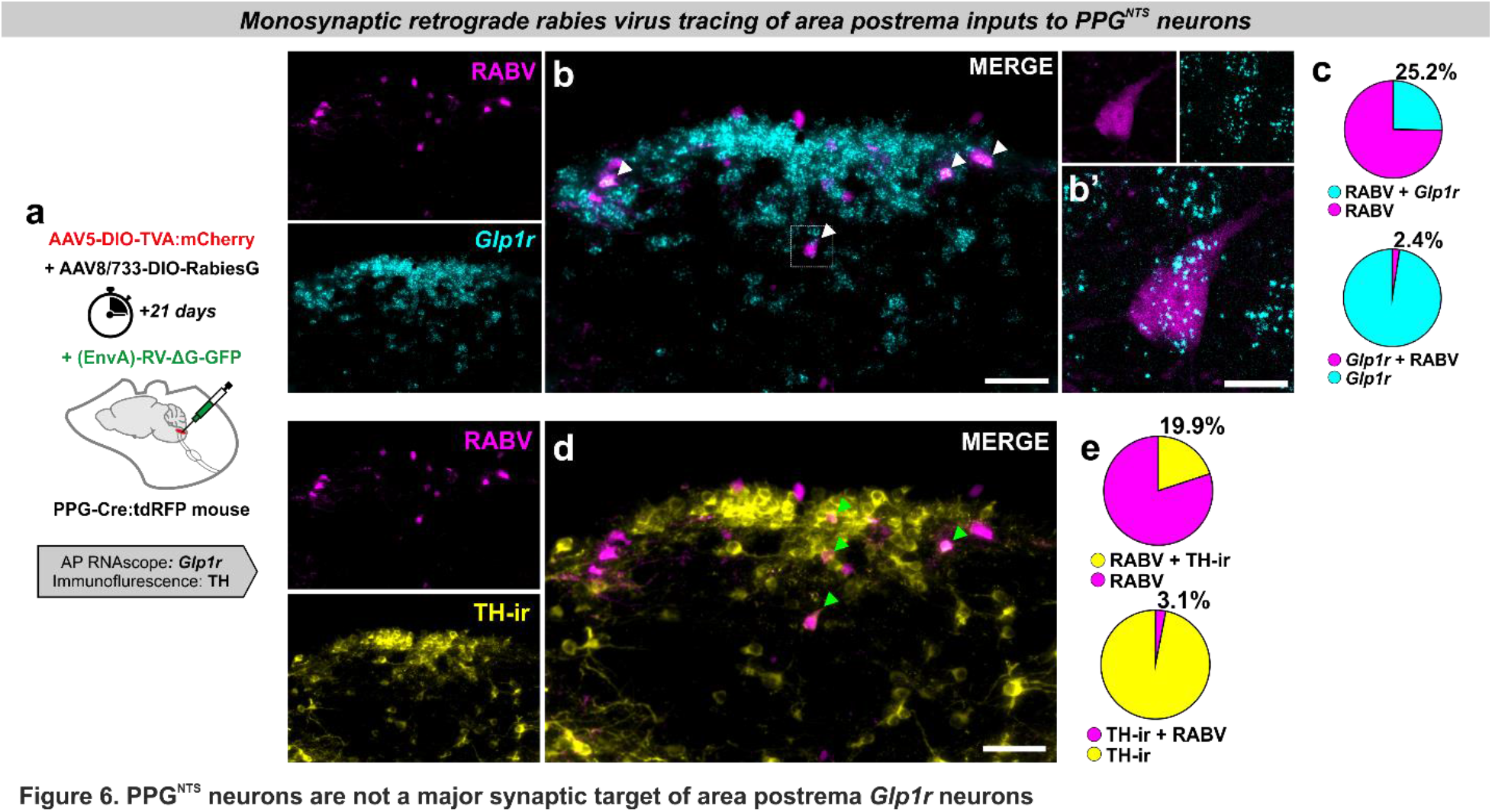
PPG^NTS^ neurons are not a major target of area postrema *Glp1r* neurons. (a) Experimental model for rabies virus-mediated monosynaptic retrograde tracing of area postrema inputs to PPG^NTS^ neurons combined with FISH for *Glp1r* (n=4 mice). (b) Photomicrographs of coronal NTS section showing RABV expression and *Glp1r* FISH. RABV+*Glp1r* co-localization shown by white arrows. Scale=100μm (inset 20μm). (c) RABV and *Glp1r* co-localization as proportions of all RABV+ cells and all *Glp1r*+ cells, from 53 RABV+ and 549 *Glp1r*+ cells. (d) Photomicrographs of coronal NTS section showing RABV expression and TH-ir. Examples of RABV+TH-ir co-localization shown by green arrows. Scale=100μm. (e) Quantification of RABV and TH-ir co-localization as proportions of all RABV+ cells and all TH-ir cells, from a total of 53 RABV+ and 341 TH-ir cells.

### Liraglutide and semaglutide suppress eating independently of PPG^NTS^ neurons

Limitations to rabies virus propagation efficiency inevitably result in an underestimate of the total number of neurons (including *Glp1r* VANs and AP neurons) providing direct synaptic input to PPG^NTS^ neurons. Nevertheless, it is apparent that the majority of *Glp1r* VANs and AP neurons are not directly presynaptic to the central GLP-1 system. However, one or both of these *Glp1r* populations may be polysynaptically connected to PPG^NTS^ neurons, in which case vagal and/or endocrine peripheral GLP-1 could still provide substantial input to the central GLP-1 system.

PPG^NTS^ neurons may alternatively (or additionally) receive input from other *Glp1r*-expressing neuronal populations which are accessible to peripheral GLP-1/GLP-1RAs and are reportedly necessary for their hypophagic effects, such as glutamatergic neurons^45^ or GABAergic NTS neurons^46^. We therefore investigated whether PPG^NTS^ neurons are a necessary component of *any* neurocircuits recruited by peripheral GLP-1RAs to suppress eating, by testing whether PPG^NTS^ neuronal ablation attenuates the anorexigenic effects of two long-acting anti-obesity GLP-1RAs, liraglutide and semaglutide (Fig. 7a). Liraglutide robustly suppressed intake and bodyweight over 24 hours in *ad libitum* eating eGFP-transduced control mice, however ablation of PPG^NTS^ neurons had no effect on the magnitude of eating suppression at any timepoint, or on 24h bodyweight loss (Fig. 7b-d, S7c-g). Semaglutide suppressed eating to an even greater extent than liraglutide, and similarly PPG^NTS^ ablation had no impact on acute or delayed eating suppression, or on bodyweight loss (Fig. 7e-g, S7h-l).

**Figure 7.**
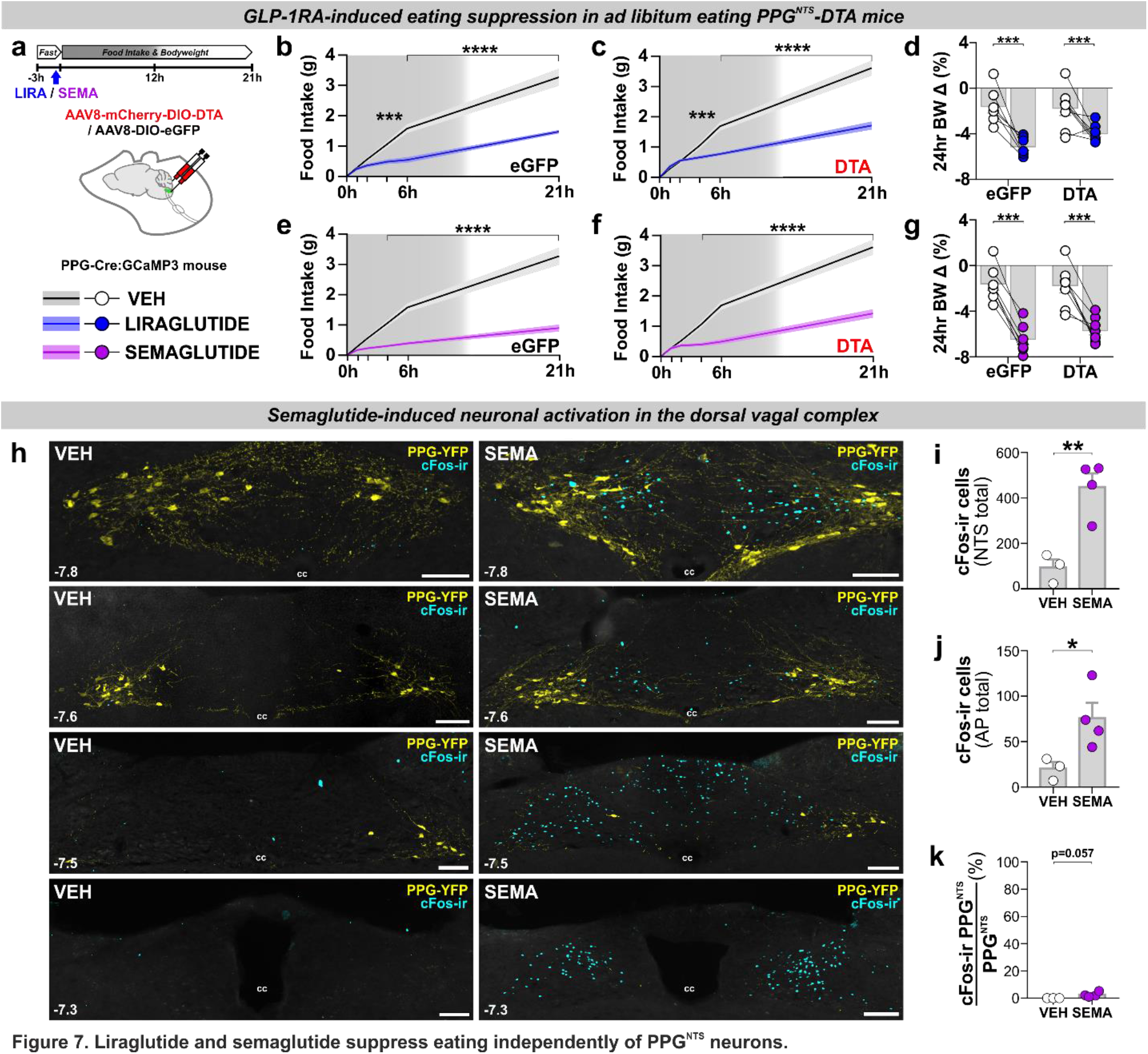
Liraglutide and semaglutide suppress eating independently of PPG^NTS^ neurons. (a) Experimental model and paradigm for GLP-1RA-induced eating suppression in PPG^NTS^-DTA ablated mice (DTA; n=8) or eGFP-transduced controls (eGFP; n=7). (b-d) Cumulative food intake and bodyweight change over 1 day in eGFP and DTA mice administered liraglutide (200 μg/kg, s.c.), 2-way within-subjects or mixed-model ANOVA: b) Drug × Time *F*_(5,30)_=35.35, *p*<0.0001; c) Drug × Time *F*_(5,35)_=74.95, *p*<0.000; d) Drug *F*_(1,13)_=33.17, *p*=0<0.0001, Virus *F*_(1,13)_=1.198, *p*=0.294. (e-g) Cumulative food intake and bodyweight change over 1 day in eGFP and DTA mice administered semaglutide (60 μg/kg, s.c.), 2-way within-subjects or mixed-model ANOVA: e) Drug × Time *F*_(5,30)_=51.83, *p*<0.0001; f) Drug × Time *F*_(5,35)_=54.28, *p*<0.0001; g) Drug *F*_(1,13)_=122.6, *p*=0<0.0001, Virus *F*_(1,13)_=0.224, *p*=0.644. (h) cFos immunoreactivity (cFos-ir) in coronal NTS sections (mm posterior to Bregma in bottom left) from PPG-YFP mice perfused 4h after vehicle (VEH) or semaglutide (SEMA; 60 μg/kg, s.c.) administration (n=7). (i-j) Total cFos in NTS and AP of mice administered vehicle (n=3) or semaglutide (n=4), unpaired 1-tailed t-tests: i) *t*_(5)_=4.59, *p*=0.0029; j) *t*_(5)_=2.66, *p*=0.0225. (k) PPG^NTS^ neurons co-localized with cFos immunoreactivity in NTS. Mann-Whitney 2-tailed U-test: U=0, *p*=0.0571.

These findings demonstrate that PPG^NTS^ neurons are not necessary for GLP-1RA-induced suppression of eating. While access to the brain by GLP-1RAs is limited, they are able to access several circumventricular *Glp1r*-expressing nuclei (in addition to the AP), which may be upstream of PPG^NTS^ neurons, or part of a subset of the *downstream* targets of these neurons^12,17,21,47^. Relevant GLP-1RA-accessible downstream *Glp1r* populations likely include neurons in the hypothalamic arcuate nucleus, which are at least partly necessary for liraglutide-induced suppression^16^. Administration of GLP-1RAs to PPG^NTS^ neuron-ablated mice cannot differentiate between whether there are any *Glp1r* populations *upstream* of PPG^NTS^ neurons that are functionally dispensable for eating suppression, or if GLP-1RAs only recruit circuits downstream or entirely independent of PPG^NTS^ neurons. We therefore investigated whether the same highly anorexigenic dose of semaglutide used in the ablation experiment was able to induce neuronal activation of PPG^NTS^ neurons, by quantifying cFos expression in the PPG-YFP mouse line^47,48^. Semaglutide induced robust cFos expression within the AP and throughout the rostro-caudal extent of the NTS (Fig. 7h-j), and additionally in hypothalamic and parabrachial nuclei (Fig. S7m-v). However, consistent with a report that the GLP-1RA Exendin-4 does not increase cFos expression in PPG^NTS^ neurons^49^, and that these neurons themselves do not express *Glp1r*^32,37^, we found that <3% of PPG^NTS^ neurons were activated by semaglutide (Figure 7k). This finding demonstrates that systemically-administered GLP-1RAs act centrally via ascending circuits parallel to, but independent of, PPG^NTS^ neurons, and/or by partially bypassing them to activate a subset of their downstream targets.

### Semaglutide and PPG^NTS^ neurons additively suppress eating

The convergent lines of neuroanatomical and functional evidence reported here suggest that, rather than comprising part of a unified GLP-1 gut-brain circuit, PPG^NTS^ neurons suppress eating via circuits which are anatomically and functionally distinct from those recruited by peripheral endogenous GLP-1 and peripherally administered GLP-1RAs. To support the hypothesis that the circuits mediating eating suppression by peripheral GLP-1RAs and PPG^NTS^ neurons are indeed entirely independent, or at least only converge at limited peripherally-accessible downstream population(s), it is a necessary to demonstrate that their concurrent activation is capable of suppressing eating in an additive manner. We therefore tested this hypothesis by administering the same dose of semaglutide that elicited robust eating suppression and neuronal activation in earlier experiments, in combination with chemogenetic activation of PPG^NTS^ neurons, and assessed intake and bodyweight over 72 hours (Fig. 8a). As expected, either manipulation alone suppressed eating over the first 24 hours, with semaglutide eliciting the stronger effect. Crucially, their combined effect was significantly additive to semaglutide alone, throughout the duration of acute chemogenetic activation (Fig. 8b-e). Consistent with our observation that PPG^NTS^ neuron activation suppresses eating without compensatory rebound hyperphagia, both cumulative intake and bodyweight were reduced at 24 and 48 hours in both of the semaglutide-treated groups. The apparent floor effect on eating suppression at 24 hours confirmed that an appropriately high dose of semaglutide was used, but this likely precluded the ability to elicit significantly additive weight loss at these later timepoints (Fig. 8f-i). Nevertheless, the additive effect of semaglutide and PPG^NTS^ activation on cumulative intake was significant even 72 hours after a single CNO dose (Fig. 8j), supporting the translational potential of this combined approach. The observed additivity could theoretically be explained by incomplete GLP-1R saturation by semaglutide within a peripherally-accessible subset of *Glp1r*-expressing nuclei downstream of PPG^NTS^ neurons. However, as we deliberately used a high dose of semaglutide to overcome this possibility, and chemogenetic activation is itself a strong and robust stimulus, this explanation is unlikely. Rather, as GLP-1RAs do not suppress eating via PPG^NTS^ neurons, and since these neurons project to numerous central nuclei involved in eating control which are not accessible to GLP-1RAs, the most parsimonious explanation is that the observed additivity must derive from concurrent activation of distinct anorexigenic neurocircuits.

**Figure 8.**
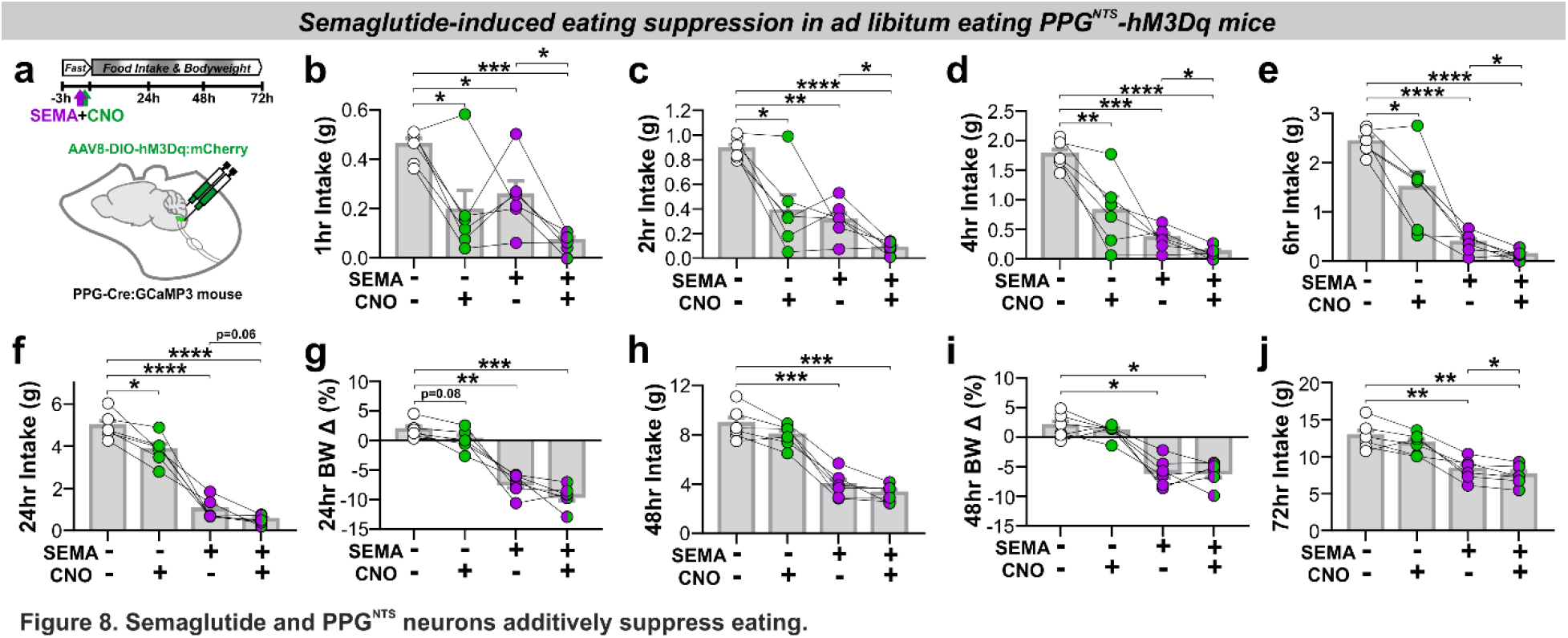
Semaglutide and PPG^NTS^ neurons additively suppress eating. (a) Experimental model and paradigm for semaglutide-induced eating suppression in PPG^NTS^-hM3Dq mice (n=6) and administered semaglutide (60 μg/kg, s.c.) and CNO (2 mg/kg, i.p.). (b-e) Cumulative food intake at 1, 2, 4 and 6 hours, 1-way within-subjects ANOVA: b) Drug *F*_(1.6,8.0)_=11.94, *p*=0.0050; c) Drug *F*_(1.5,7.7)_=22.12, *p*=0.0009; d) Drug *F*_(1.3,6.3)_=35.35, *p*=0.0006; e) Drug *F*_(1.2,6.0)_=40.72, *p*=0.0005. (f-j) Cumulative food intake and bodyweight change at 24, 48 and 72 hours, 1-way within-subjects ANOVA: f) Drug *F*_(1.6,8.2)_=125.8, *p*<0.0001; g) Drug *F*_(2.1,10.5)_=61.61, *p*<0.0001; h) Drug *F*_(2.3,11.3)_=102.7, *p*<0.0001; i) Drug *F*_(2.1,10.6)_=24.38, *p*<0.0001; j) Drug *F*_(1.9,9.3)_=40.35, *p*<0.0001. 72hr BW data not shown: Drug *F*_(2.0,10.2)_=4.22, *p*=0.0454, no significant pairwise comparisons.

## Discussion

Here we report that PPG^NTS^ neurons encode satiation specifically during large meals, and have capacity for pharmacological activation to suppress eating without compensatory rebound hyperphagia or behavioural disruption. Activation of *Glp1r* VANs similarly suppressed intake, but did condition flavour avoidance, and complementary circuit mapping approaches demonstrated that PPG^NTS^ neurons are not a major synaptic target of this vagal population. We report that PPG^NTS^ neurons instead predominantly receive vagal input from *Oxtr* VANs, and are required for peripheral oxytocin-induced eating suppression. Similarly, PPG^NTS^ neurons are at most a minor synaptic target of *Glp1r* neurons in the area postrema, suggesting that endocrine GLP-1 signalling from the periphery by this route does not require PPG^NTS^ neurons. Consistent with this observation, PPG^NTS^ neurons are not recruited by peripherally administered semaglutide, or required for the anorexigenic effects of liraglutide or semaglutide, and concurrent administration of semaglutide and activation of PPG^NTS^ neurons suppresses eating in an additive manner. We therefore conclude that, contrary to the prevalent hypothesis, the peripheral and central GLP-1 systems are components of anatomically and functionally independent eating control circuits. Furthermore, while pharmacokinetically-optimized GLP-1RAs may access a limited subset of *Glp1r* neuron populations downstream of PPG^NTS^ neurons, such partial convergence of recruited circuits does not preclude additive suppression of eating. PPG^NTS^ neurons are thus a rational pharmacological target for obesity treatment both in their own right and for combination therapy with GLP-1RAs.

## Acknowledgements

We would like to thank Lotte Bjerre Knudsen at Novo Nordisk for valuable discussions and provision of liraglutide and semaglutide. We would also like to thank Myrtha Arnold at ETH Zurich and Yalun Tan at the University of Florida for their expert technical assistance, and Kevin Beier at UC Irvine for providing rabies virus for retrograde tracing. This study was supported by MRC project grant MR/N02589X/1 to ST. The initial collaboration between the Trapp, de Lartigue and Langhans labs which generated this work was made possible thanks to a UCL Global Engagement Fund award to DB, and a UCL Neuroscience ZNZ Collaboration award to ST and WL. Research in the de Lartigue lab was funded by the NIH (NIDDK grant R01 DK116004) and with institutional support from the University of Florida College of Pharmacy. Research in the Reimann/Gribble laboratories was funded by the Wellcome Trust (106262/Z/14/Z and 106263/Z/14/Z) and the MRC (MRC_MC_UU_12012/3). Research in the Rinaman laboratory was funded by the U.S. National Institutes of Health (MH059911 and DK100685).

## Author Contributions

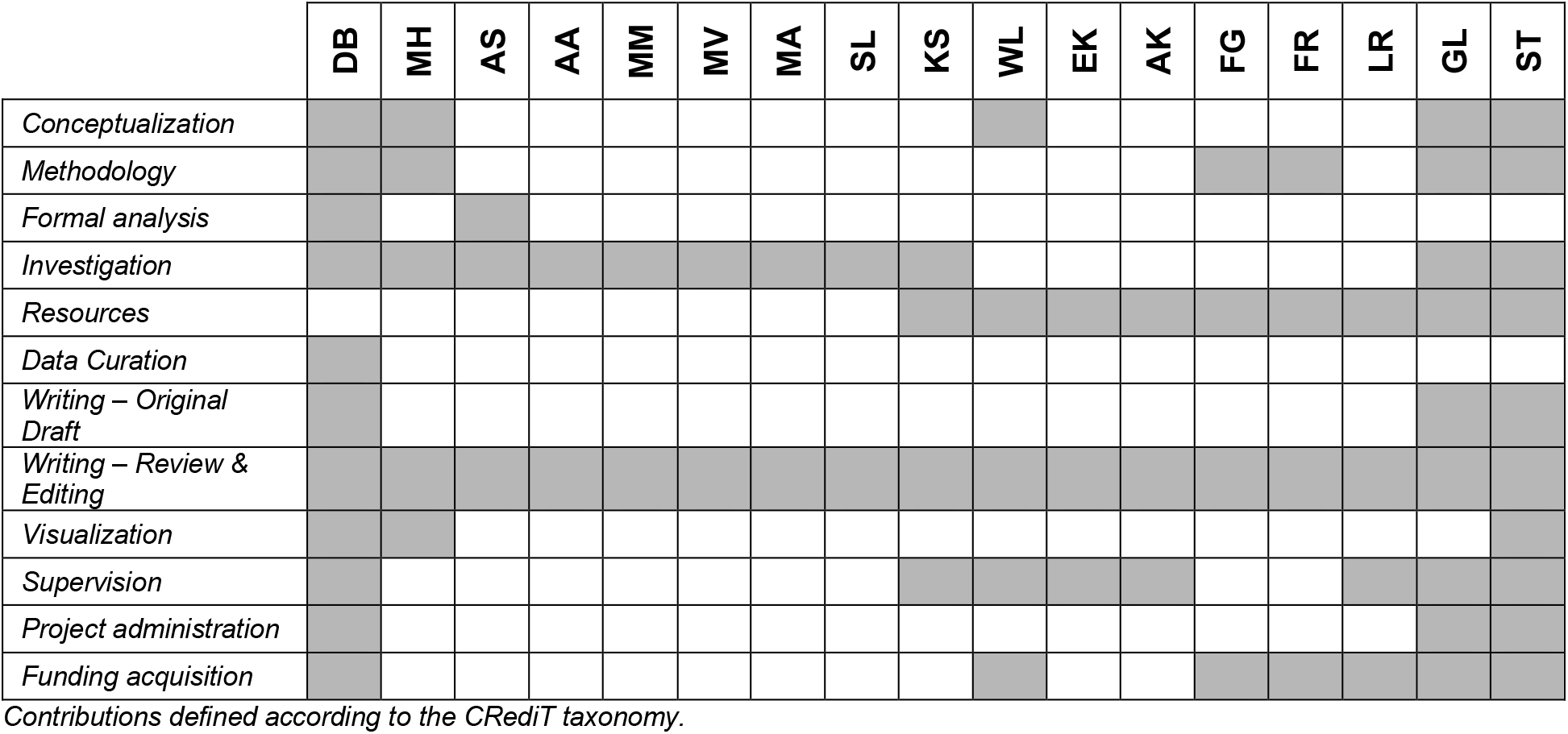

## Competing interests statement

The FR + FMG laboratory receives funding from AstraZeneca, Eli Lilly and LGM for unrelated research and FMG consults for Kallyope (New York). All other authors have nothing to declare.

## Methods

### Animals

We used 116 mice of both sexes (≥10 weeks) from five previously reported strains, all maintained on C57BL/6 backgrounds and bred in house. For selective Cre-dependent viral targeting and *ex vivo* Ca^2+^ imaging of PPG neurons, we used mGlu-Cre/tdRFP ^50^ and mGlu-Cre/GCaMP3 strains ^38^, referred to herein as PPG-Cre:tdRFP and PPG-Cre:GCaMP3, respectively. For visualization of PPG neuron somata, axons and dendrites, we used the mGlu-YFP strain (Reimann et al., 2008; referred to as PPG-YFP). For selective Cre-dependent viral targeting of GLP-1 receptor-expressing neurons we used the *Glp1r*-Cre/tdRFP strain ^51^, or a cross with the PPG-YFP strain (Card et al., 2018; referred to as GLP-1R-Cre x PPG-YFP). All mice were kept on a 12h light/dark cycle with chow and water available *ad libitum* and group housed until surgery and/or behavioural experiments. Within-subjects design experiments were conducted using sex-balanced cohorts of appropriate genotype littermates as far as possible. Similarly, for between-subjects and mixed model design experiments, littermates were semi-randomly allocated to virus groups to ensure groups were balanced for sex and age as far as possible. Power calculations were not performed, appropriate group sizes (detailed in figure legends) were determined from pilot experiments and our previously published studies using these models and behavioural paradigms ^7,49,52^.

Experiments conducted in the UK were performed in accordance with the U.K. Animals (Scientific Procedures) Act 1986 and with appropriate institutional approval. Experiments conducted in the U.S. were performed in accordance with the U.S. Public Health Service’s Policy on the Humane Care and Use of Laboratory Animals and experimental protocols were approved by the Institutional Animal Care and Use Committees of Florida State University and the University of Florida. Experiments conducted in Switzerland were approved by the Canton of Zurich’s Veterinary Office (ETH Zurich).

### Stereotaxic Surgery

#### NTS Virus Injections

Injection of viral vectors targeted to the caudal NTS were performed as previously described ^7,26,49^. Mice were anaesthetized with intramuscular medetomidine (1 mg/ kg) + ketamine hydrochloride (50 mg/kg) or 1.5-2.5% isoflurane, and given carprofen analgesia (5mg/kg, s.c.). Core temperature was maintained using a homeothermic monitoring system, and appropriate depth of surgical anaesthesia was determined by absence of pedal reflex. The skull was restrained in a stereotaxic frame, and the head flexed downwards such that the nose and neck were at a right angle. The scalp was incised from the occipital crest to first vertebrae, and muscle layers parted to expose the atlanto-occipital membrane. The membrane was bisected horizontally with a 30G needle to expose the brainstem surface with obex as an anatomical landmark. Viral vectors (as detailed in figures and Methods Table 1) were injected via pulled glass micropipettes at the following coordinates from obex: +0.1mm rostral, ±0.5mm lateral and −0.35mm ventral. Viruses encoding chemogenetic effectors, diptheria toxin subunit A (DTA) and control reporters were all bilaterally injected in volumes of 200-250nl. Mice were allowed to recover for a minimum of 3 weeks before behavioural experiments began. For monosynaptic retrograde tracing from PPG neurons in the caudal NTS, 300nl of a 1:1 mix of AAV5-EF1a-FLEX-TVA:mCherry and AAV8/733-CAG-FLEX-RabiesG were bilaterally injected +0.1mm rostral, ±0.4mm lateral and 0.35mm ventral to obex. 21 days later, 400nl in total of (EnvA)-RABV-∆G-GFP was unilaterally injected at two injection sites: +0.1mm rostral, +0.25mm lateral and 0.35-0.45mm ventral to obex; and +0.1mm rostral, +0.4mm lateral and 0.35-0.45mm ventral to obex. Mice were transcardially perfused for histological processing and in situ hybridisation 7 days later. We have previously optimised, validated and published all viral targeting strategies used in the present study ^7,26,49,52^. Post-mortem tissue sections were processed from all mice used in behavioural studies and verified for appropriate expression of fluorescent reporters. Mice in which viral injection targeting was inaccurate, or transduction efficiency below expected levels, were omitted from analyses.

**Table 1.**
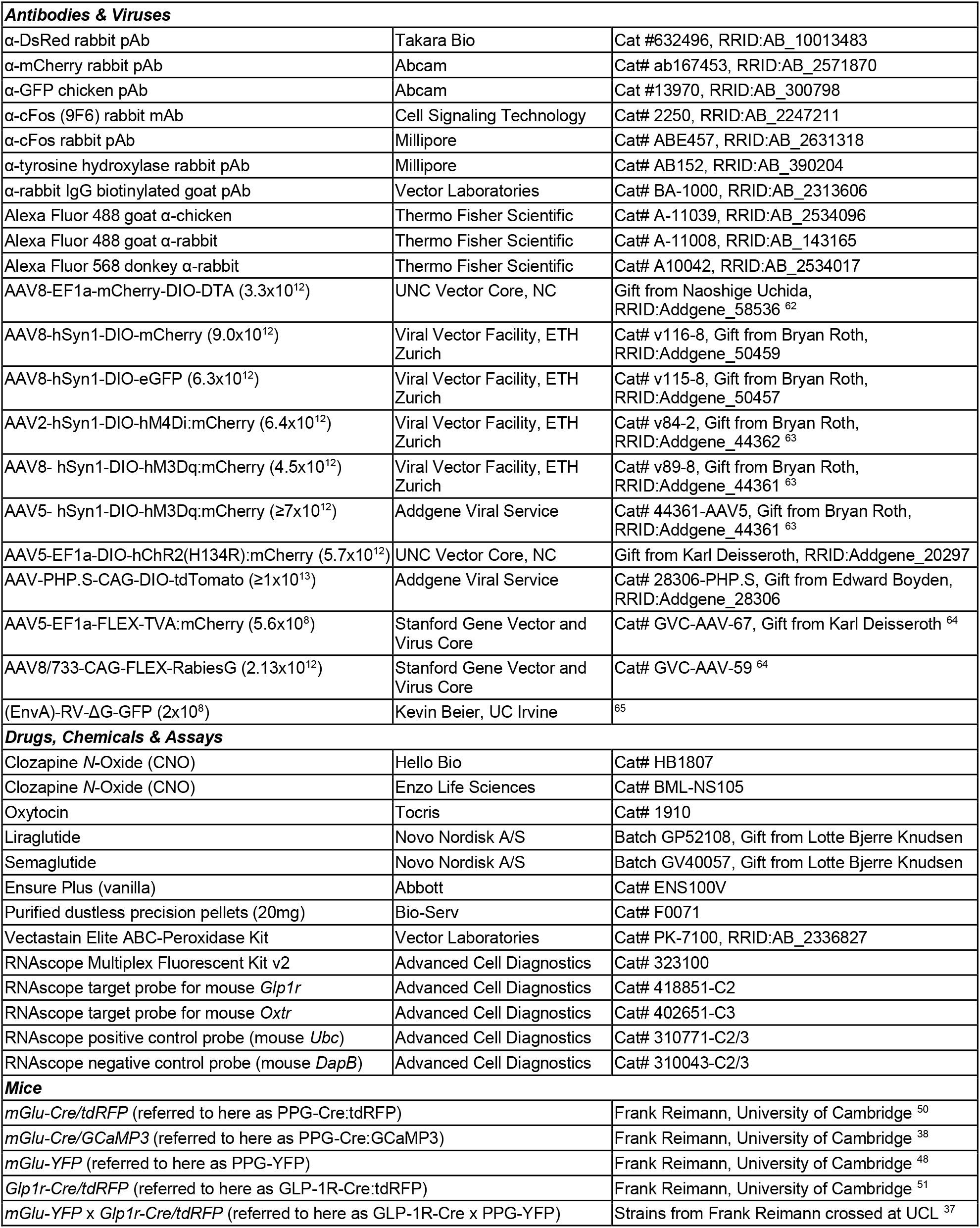
Reagents and resources details.

#### Nodose Ganglia Virus Injections

Injection of viral vectors targeted to the somata of vagal afferent neurons in the nodose ganglia were performed as previously described ^52^. Mice were anaesthetized with 1.5-2.5% isoflurane and given carprofen analgesia (5mg/kg, s.c.), then the ventral surface of the neck was incised, and muscles parted to expose the trachea. The vagus nerve was separated from the carotid artery to allow access to the nodose ganglia. In each nodose ganglia, a total volume of 500nl of viral vector (AAV5-hSyn1-DIO-hM3Dq:mCherry, AAV5-EF1a-DIO-hChR2(H134R):mCherry or AAV-PHP.S-CAG-DIO-tdTomato, as detailed in figures) was injected into sites rostral and caudal to the laryngeal nerve branch, using a bevelled tip pulled glass micropipette and pneumatic microinjector. Viruses encoding chemogenetic and optogenetic effectors were injected bilaterally, and the virus for tdTomato-visualized projection tracing was injected unilaterally into left or right nodose ganglia. Mice were allowed to recover for a minimum of 2 weeks before behavioural experiments or transcardial perfusion.

#### Optical Fibre Implantation

Two weeks after bilateral injection of AAV5-EF1a-DIO-hChR2(H134R):mCherry in nodose ganglia, optical fibres were prepared and implanted as previously described ^53^. Optical fibres (CFLC230–10 ceramic ferrules with FT200UMT fibre, Thorlabs) were unilaterally implanted above the right caudal NTS, 7.5mm caudal, 0.25mm lateral and 4.0mm ventral to Bregma. Skull screws, superglue and dental cement were used to secure the fibre, and mice were allowed to recover for an additional 2 weeks prior to behavioural testing.

### Behavioural Studies

#### Drug Administration

Clozapine *N*-oxide (CNO; Hello Bio / Enzo) was administered intraperitoneally at 2 mg/kg in 2 ml/kg dose volume for all experiments, typically 15 minutes prior to dark onset. We have previous determined that in our hands CNO at this dose does not affect eating behaviours in control virus transduced mice, and similarly that the chemogenetic effectors used do not have any constitutive activity which affects eating behaviours ^7,52^. To minimize animal use and maximize statistical power, all chemogenetic experiments in the present study were therefore conducted in hM3Dq-expressing mice using a within-subjects design. All mice received both CNO and saline vehicle in a counterbalanced manner, and hence acted as their own controls. Similarly, when assessing the effect of PPG^NTS^ ablation on the anorexigenic actions of oxytocin, liraglutide and semaglutide, we used a mixed model design, whereby mice in DTA-ablated and control cohorts all received drug and vehicle in a counterbalanced manner. Oxytocin (Tocris) was administered intraperitoneally 15 minutes prior to dark onset at 0.4 mg/kg, based on reports that this dose and route of administration elicits vagal-dependent eating suppression in mice ^40,41^. Liraglutide and semaglutide (gift from Lotte Bjerre Knudsen, Novo Nordisk) were administered subcutaneously 30 minutes prior to dark onset at 0.2 mg/kg and 0.06 mg/kg, respectively, in 5 ml/kg dose volume, based on recommendations from LBJ and previous reports of the anorexigenic effects of these drugs in mice ^16,17,54^.

#### Eating Behaviour Paradigms

Drug- or neuronal manipulation-induced changes to eating behaviour were assessed from dark onset in *ad libitum* eating or fasted mice. In the *ad libitum* eating paradigm, mice were habituated (≥5 sessions) to being fasted for the final 3 hours of the light phase and receiving saline/vehicle injections 5-30 mins prior to return of food at dark onset. This protocol minimized hypophagia from handling and injection stress, and entrained mice to eat consistently from dark onset, without needing to induce negative energy balance. All experiments using either manual or automated measurement of food intake used this protocol for assessment of *ad libitum* eating, except for metabolic phenotyping of PPG^NTS^ ablated mice experiment, in which mice were not handled or injected and were already habituated to test cages. To assess the effect of chemogenetic manipulations on the behavioural satiety sequence, and large meals driven by refeeding after a prolonged fast, mice were fasted for 18 hours prior to dark onset. To assess the effect of optogenetic activation of *Glp1r*-expressing vagal afferent neurons on acute feeding, mice were fasted for the entire light phase and intake measured during the first 30 minutes of the dark phase. The effect of optogenetic activation was also assessed using a within-subjects design, with all mice tested for 30 minute intake under ‘laser on’ (20ms blue light pulse every 3 seconds, ~5mW intensity) and ‘laser off’ (tethered but no light pulses) conditions in a counterbalanced manner.

#### Food Intake Measurement

Food intake was measured manually, using open source FED pellet dispensers (Feeding Experimentation Device; Nguyen et al., 2016), or using commercially-available Phenomaster (TSE Systems) or Promethion (Sable Systems) systems. For manual measurement of intake in the *ad libitum* eating paradigm, mice were weighed at the start of the 3 hour fast, then a pre-weighed amount of food was returned at dark onset. Food was again weighed at 1, 2, 4, 6 (GLP-1RA experiments only), and 21 hours, at which point 24 hour bodyweight change was also determined. FED dispensers were used for all experiments involving chemogenetic manipulations of PPG^NTS^ neurons, the Phenomaster system for chemogenetic activation of *Glp1r*-expressing vagal afferent neurons, and the Promethion system for metabolic phenotyping of PPG^NTS^ ablated mice. For all food intake measurement systems and eating behaviour paradigms, mice were habituated to the test equipment and all aspects of the paradigm (including vehicle dosing where appropriate) prior to the start of testing. Mice were considered habituated after their intakes during ≥3 consecutive habituation sessions were not significantly different, and no directional trend was apparent. Meal pattern analysis was conducted on automated food intake data from experiments testing the effect of inhibition or ablation of PPG^NTS^ neurons. A meal was defined as the sum of all bouts ≥0.02g with intra-meal intervals <10 minutes, based on the standard operating procedure of the UC Davis Mouse Metabolic Phenotyping Centre ^55^. Consumption of Ensure liquid diet was measured manually, and the temporal pattern of Ensure drinking was measured by offline video coding of licking duration (blinded to drug treatment), using the BORIS open source video coding software package ^56^.

#### Behavioural Satiety Sequence Analysis

Alterations to the behavioural satiety sequence (BSS) following chemogenetic activation of PPG^NTS^ neurons were determined using the continuous monitoring BSS protocol ^57,58^, adapted for use with mice and FED dispensers. Mice were fasted for 18 hours, FED dispensers were returned to home cages at dark onset, and behaviour recorded for 40 minutes using infrared video cameras. Behaviours were subsequently coded offline using BORIS software by trained observers (Cohen’s κ for inter-rater reliability >0.9) blinded to treatment group. Behaviours were coded as mutually-exclusive categories: eating, drinking, grooming (including scratching), inactive (resting and sleeping) and active (locomotion and rearing). In pilot experiments, the duration of water drinking was found to be extremely low and unaffected by our manipulations, hence was omitted from further analyses. The total duration mice spent exhibiting behaviours in the remaining 4 categories were calculated for 8 × 5 minute time bins. For qualitative and semi-quantitative evaluation of the stochastic sequence of satiety behaviours (eating → grooming → inactive), the mean durations of these 3 categories were plotted across all time bins separately for saline control and CNO activated conditions. To aid visualization of neuronal activation-induced acceleration of the sequence, horizontal dotted lines were added denoting the time bin during which the (probabilistic) transition from feeding to resting occurs, which is typically considered the satiation point / onset of satiety. Data were also presented and analysed to quantitatively test the effect of chemogenetic activation over time in each behaviour category.

#### Indirect Calorimetry

We measured respiratory exchange ratio and energy expenditure concurrently with food intake from PPG^NTS^ ablated and *Glp1r*^Nodose^-hM3Dq mice, using the indirect calorimetry functionality integrated into the Phenomaster and Promethion systems. Mice were habituated to test cages for ≥3 days before testing, and metabolic data were collected for 24 hours for between-subjects analysis (PPG^NTS^-DTA ablated vs PPG^NTS^-mCherry controls), or during two 24 hour test sessions (*Glp1r*^Nodose^-hM3Dq, counterbalanced for CNO administration), separated by ≥48 hour washout periods during which mice remained in test cages. Respiratory exchange ratio (RER) was obtained from measurement of mice’s O2 consumption (ml/kg/hr) and CO2 production (ml/kg/hr), using the equation: RER = VCO2 / VO2. Energy expenditure (EE) was calculated using the Weir equation: EE = 3.941 × VO2 + 1.106 × VCO2. Raw data from both systems were used to generate standardized output files, which were imported into the CalR analysis tool ^59^ for production and analysis of intake and metabolic data over light and dark phases and total circadian cycle. As now recommended for analysis of calorimetry data from these systems, energy expenditure was not normalized to bodyweight.

#### Conditioned Flavour Preference

Whether optogenetic activation of *Glp1r* vagal afferent neurons conditioned a preference for (or avoidance of) a flavour was assessed as previously described ^52^. Experiments were conducted within sound-attenuated cubicles, using behavioural chambers equipped with two sipper tubes connected to contact-based licking detection devices, allowing high resolution measurement of licking responses (Med Associates Inc.). Following recovery from surgeries for nodose ganglia virus injection and optical fibre implantation, individually-housed mice were placed on a food and water restriction regime, under which they were maintained at 90% of starting bodyweight and were limited to 6 hours of water access per day. Mice were habituated to behavioural chambers (including being tethered to fibre cables) and trained to lick for a 0.025% saccharin solution during daily 1 hour habituation sessions, conducted during the light phase. Mice were considered trained to saccharin licking once they showed <10% between-session variability in the number of licks, a criterion all mice reached within 10 sessions.

Once trained, a ‘pre’-test was conducted in which mice were given access to two novel Kool-Aid flavours (cherry or grape, both 0.05% in 0.025% saccharin solution) for 10 minutes, with sipper bottle positions switched after 5 minutes to avoid position bias. Mice then underwent 3 × 1 hour training sessions for each flavour (alternately over 6 days), in which both bottles contained the same flavour. One flavour was paired with laser stimulation (CS+), such that licking triggered blue light laser stimulation via a TTL output signal. Specifically, 10 licks triggered a 20ms light pulse of ~5mW intensity, with additional licks during the following 10 seconds having no programmed consequences. Further bouts of ≥10 licks triggered additional pulses in the same manner throughout the 1 hour session. During training sessions with the unpaired (CS-) flavour, mice were tethered but licking did not elicit laser stimulation. Upon completion of these training sessions, mice underwent a ‘post’-test identical to the ‘pre’-test, i.e. both flavours were available, and licking did not elicit laser stimulation. The number of licks for the laser-paired flavour during ‘pre’ and ‘post’ tests was used to calculate preference ratios (CS+ licks / total licks) for the flavour before and after training, to determine if optogenetic stimulation of *Glp1r* vagal afferent neurons increased or decreased preference for the paired flavour.

### Immunohistochemistry & In Situ Hybridization

#### Tissue Preparation

Mice were deeply anaesthetized then transcardially perfused with ice-cold PB/PBS (0.1M, pH 7.2) then 4% formaldehyde in PB/PBS. Brains and NG (when required) were extracted and post-fixed in 4% formaldehyde at 4°C overnight (≤2 hours for NG), before being cryoprotected in 20-30% sucrose solution for ≥24 hours at 4°C. Brains were sectioned into 30-35μm coronal sections, collected free-floating and stored at 4°C until processing for immunofluorescent labelling as detailed below. NG were sectioned into 10μm sections, collected on Superfrost Plus microscope slides and stored at −20°C until processing for *in situ* hybridization as detailed below.

#### Immunofluorescent labelling

Brain sections were processed for amplification of fluorescent reporter signals by immunofluorescent labelling of tdRFP, mCherry, eYFP, eGFP and/or GCaMP3 as previously described ^7^. Briefly, sections were incubated free-floating with primary antibodies (see Methods Table 1) overnight at 4°C in PBS with 2% normal goat/donkey serum and 1% BSA, followed by 2 hours at room temperature with secondary antibodies conjugated to fluorophores appropriate for the native fluorescent reporter being amplified (i.e. Alexa Fluor 488 for eYFP/eGFP/GCaMP3 and Alexa Fluor 568 for tdRFP/mCherry).

#### RNAscope *In situ* Hybridization (Nodose Ganglia)

Sections from nodose ganglia of mice previously injected with viruses for monosynaptic retrograde rabies tracing were processed for *in situ* hybridization of *Glp1r* and *Oxtr* mRNA using the RNAscope assay as previously reported ^60^. Sections were cut at 10μm on a cryostat and collected on Superfrost Plus slides, then allowed air-dry at room temperature for one hour.

Slides were then dipped in molecular grade ethanol and further air-dried overnight at room-temperature. RNAscope *in situ* hybridization was performed on these sections using the RNAscope Multiplex Fluorescent Kit v2 (Advanced Cell Diagnostics) as per the manufacturer’s instructions, with a modification to the pre-treatment procedure (Protease IV incubation conducted for 20 min at room temperature) that allows for preservation of the fluorescent reporter signal while also providing optimal signal from the target mRNAs. Probes for *Glp1r*, *Oxtr* and appropriate positive (*Ubc*) and negative (*DapB*) controls (detailed in Methods Table 1) were hybridized and after completion of the procedure slides were immediately cover slipped using Prolong Antifade medium.

#### RNAscope *In situ* Hybridization (Brainstem)

Brainstem sections containing the area postrema were pre-treated with hydrogen peroxide for 30 minutes at room temperature, slide-mounted in dH_2_O and air dried overnight. Sections were subsequently processed for *in situ* hybridization of *Glp1r* mRNA using the same reagents and protocol as nodose ganglia sections, followed by additional processing for immunofluorescent labelling of GFP and tyrosine hydroxylase (TH). Incubation with primary antibodies against GFP and TH was performed concurrently overnight at room temperature, with the remaining protocol conducted as described above for labelling of fluorescent reporters. Sections were then dehydrated in increasing concentrations of ethanol, cleared in xylene and cover slipped using Cytoseal 60.

#### cFos Quantification

For immunohistochemical validation of chemogenetic PPG^NTS^ neuron activation, brains from PPG^NTS^-hM3Dq mice were processed for immunofluorescent labelling of cFos as previously reported ^7^. For quantification of semaglutide-induced cFos expression, mice expressing eYFP in PPG neurons were habituated to handling and the standard *ad libitum* eating behaviour paradigm, including vehicle injection 30 minutes prior to dark onset. Food intake manually quantified after 3 hours during habituation sessions and on the day semaglutide was administered, allowing a within-subjects quality control for the effect of semaglutide in this experiment. On the test day, mice were injected with vehicle or semaglutide (0.06mg/kg as per behavioural studies) and transcardially perfused 4 hours later. Coronal brain sections were prepared as above, then processed for immunoperoxidase labelling of cFos with DAB-Ni followed by immunofluorescent amplification of eYFP as previously described ^26^.

#### Imaging

Brain sections labelled for fluorescent reporters and/or cFos expression were imaged using an upright epifluorescence and brightfield microscope (Leica) with a Retiga 3000 CCD camera (QImaging). For co-localization of DAB-Ni labelled cFos and PPG-eYFP neurons, brightfield and fluorescence images were sequentially captured in the same focal plane. Quantification of cFos expression and co-localization was conducted using merged native brightfield (DAB) and fluorescent (PPG-eYFP) images. For clarity of presentation, brightfield DAB images were inverted and pseudocolored prior to merging with fluorescent channels. Nodose ganglia and brainstem sections processed for *in situ* hybridization and/or fluorescent reporters were imaged with a Keyence BZ-x700 at 20x or 40x in 0.6μm optical sections, or a Leica TCS SP8 confocal microscope at 20x. For imaging of sections processed for *in situ* hybridization, sections hybridized with positive and negative control probes were used to determine exposure time and image processing parameters necessary for optimal visualization of mRNA signals and control for possible degradation. Generation of montages from individual images, brightness and contrast adjustment, and quantification of cFos expression using the Cell Counter plugin were all performed using Fiji open source biological image analysis software ^61^.

### Brain Slice Ca^2+^ Imaging

#### Imaging Data Capture

Coronal brainstem slices (200μm) were obtained from PPG-Cre:GCaMP3 mice and used to assess the effects of bath-applied oxytocin on PPG^NTS^ neuron calcium dynamics using methods previously described in detail ^39^. Oxytocin was dissolved in aCSF (3mM KCl, 118mM NaCl, 25mM NaHCO_3_, 5mM glucose, 1mM MgCl_2_, 2mM CaCl_2_; pH 7.4) to give a bath concentration of 100nM, based on reports that this concentration elicits robust activation of vagal afferent neurons under *ex vivo* conditions ^40^. Slices were superfused with aCSF for ≥10 minutes, with the final 5 minute period prior to oxytocin application used to determine baseline fluorescence intensity. Slices were then superfused with oxytocin solution for 3-5 minutes, washed with aCSF for ≥10 minutes, then finally superfused with 100μM glutamate for 1 minute as a positive control to confirm imaged neurons were healthy and responsive to glutamatergic input. GCaMP3 fluorescence was excited at 460 ± 25 nM using an LED light source, for 250ms every 5 seconds. Imaging was conducted using a widefield microscope (Zeiss) with 40x water immersion lens and captured at 12-bit on a CCD camera (QClick, QImaging). Data were obtained from 8 experiments (i.e. recordings from single slices) from 3 mice.

#### Imaging Data Analysis

Time-lapse image recordings were imported into FIJI software, with the StackReg plugin used to correct for XY drift. Regions of interest (ROIs) were manually drawn around all PPG^NTS^ somata in the field of view, with additional ROIs used to determine background intensity for each experiment. Background intensity was subtracted from ROIs and a cubic polynomial function was used to adjust for bleaching. Data are presented as Δ*F*/*F*_0_, where *F*_0_ is the mean fluorescence intensity over the 5 minute baseline period, and Δ*F* is the intensity at each timepoint with *F*_0_ subtracted. Response magnitudes were determined using the area under the curve (AUC) over 4 minutes from first application of oxytocin. To ensure artifactual fluctuations were not included in analyses, only fluorescence changes for which the magnitude was greater (or less) than 3 standard deviations of the baseline period AUC were considered to be ‘responders’. As the noise level (i.e. variability in baseline AUC) differs between slice recordings, this threshold is not absolute, hence there is some degree of overlap between the AUC of ‘non-responsive’ ROIs from more noisy recordings and ‘responsive’ ROIs from less noisy recordings. As a further quality control, oxytocin-responsive ROIs were only included for analysis if they subsequently were responsive to glutamate.

### Quantification and statistical methods

Data are presented as mean ± SEM, and were analysed for statistical significance as detailed in figure legends using Student’s *t*-test, one-way within-subjects or two-way within-subjects/mixed-model ANOVA (with the Greenhouse-Geisser correction applied where appropriate). Where data were not normally distributed, non-parametric equivalents were used as detailed.

Significant one-way ANOVA tests were followed by pairwise comparisons with Tukey’s correction for multiple comparisons. For two-way ANOVA, either simple main effects were reported, or significant interactions were reported and followed by pairwise comparisons with Sidak’s correction for multiple comparisons. The threshold for statistical significance was considered <0.05, and significant comparisons are reported in all figures as: * p<0.05, ** p<0.01, *** p<0.001, **** p<0.0001. For transparency, all comparisons in which p<0.1 (but ≥0.05) are additionally reported with exact p values shown. Statistical analyses were conducted using GraphPad Prism 7 or IBM SPSS Statistics 26.

### Data availability

The datasets supporting the current study are available from the lead contact on request. All mouse lines, plasmids and reagents used in this study have been previously published and/or are commercially available. Further information and requests for resources and reagents should be directed to and will be fulfilled by Prof. Stefan Trapp.

## References

1. Knudsen, L. B. & Lau, J. The discovery and development of liraglutide and semaglutide. Front. Endocrinol. (Lausanne). 10, 155 (2019).

2. Turton, M. D. et al. A role for glucagon-like peptide-1 in the central regulation of feeding. Nature 379, 69–72 (1996).

3. Trapp, S. & Richards, J. E. The gut hormone glucagon-like peptide-1 produced in brain: is this physiologically relevant? Curr. Opin. Pharmacol. 13, 964–969 (2013).

4. Müller, T. D. et al. Glucagon-Like Peptide 1 (GLP-1). Mol. Metab. 30, 72–130 (2019).

5. Gaykema, R. P. et al. Activation of murine pre-proglucagon–producing neurons reduces food intake and body weight. J. Clin. Invest. 127, 1031–1045 (2017).

6. Liu, J. et al. Enhanced AMPA Receptor Trafficking Mediates the Anorexigenic Effect of Endogenous Glucagon-like Peptide-1 in the Paraventricular Hypothalamus. Neuron 96, 897–909.e5 (2017).

7. Holt, M. K. et al. Preproglucagon neurons in the nucleus of the solitary tract are the main source of brain GLP-1, mediate stress-induced hypophagia, and limit unusually large intakes of food. Diabetes 68, 21–33 (2019).

8. Zheng, H., Stornetta, R. L., Agassandian, K. & Rinaman, L. Glutamatergic phenotype of glucagon-like peptide 1 neurons in the caudal nucleus of the solitary tract in rats. Brain Struct. Funct. 220, 3011–3022 (2015).

9. Trapp, S. et al. PPG neurons of the lower brain stem and their role in brain GLP-1 receptor activation. Am. J. Physiol. Regul. Integr. Comp. Physiol. 309, R795–804 (2015).

10. Cheng, W. et al. Leptin receptor–expressing nucleus tractus solitarius neurons suppress food intake independently of GLP1 in mice. JCI Insight 5, e134359 (2020).

11. Merchenthaler, I., Lane, M. & Shughrue, P. Distribution of pre-pro-glucagon and glucagon-like peptide-1 receptor messenger RNAs in the rat central nervous system. J. Comp. Neurol. 403, 261–80 (1999).

12. Cork, S. C. et al. Distribution and characterisation of Glucagon-like peptide-1 receptor expressing cells in the mouse brain. Mol. Metab. 4, 718–731 (2015).

13. Drucker, D. J. Mechanisms of Action and Therapeutic Application of Glucagon-like Peptide-1. Cell Metab. 27, 740–756 (2018).

14. Krieger, J.-P. Intestinal glucagon-like peptide-1 effects on food intake: physiological relevance and emerging mechanisms. Peptides 170342 (2020). doi:10.1016/j.peptides.2020.170342

15. Grill, H. J. A role for GLP-1 in treating hyperphagia and obesity. Endocrinology bqaa093 (2020). doi:10.1210/endocr/bqaa093

16. Secher, A. et al. The arcuate nucleus mediates GLP-1 receptor agonist liraglutide-dependent weight loss. J. Clin. Invest. 124, 4473–88 (2014).

17. Gabery, S. et al. Semaglutide lowers body weight in rodents via distributed neural pathways. JCI Insight 5, e133429 (2020).

18. Varin, E. M. et al. Distinct Neural Sites of GLP-1R Expression Mediate Physiological versus Pharmacological Control of Incretin Action. Cell Rep. 27, 3371–3384.e3 (2019).

19. Kanoski, S. E., Fortin, S. M., Arnold, M., Grill, H. J. & Hayes, M. R. Peripheral and central GLP-1 receptor populations mediate the anorectic effects of peripherally administered GLP-1 receptor agonists, liraglutide and exendin-4. Endocrinology 152, 3103–3112 (2011).

20. Punjabi, M. et al. Circulating Glucagon-like Peptide-1 (GLP-1) Inhibits Eating in Male Rats by Acting in the Hindbrain and Without Inducing Avoidance. Endocrinology 155, 1690–1699 (2014).

21. Ast, J. et al. Super-resolution microscopy compatible fluorescent probes reveal endogenous glucagon-like peptide-1 receptor distribution and dynamics. Nat. Commun. 11, 1–18 (2020).

22. Krieger, J. P. et al. Knockdown of GLP-1 receptors in vagal afferents affects normal food intake and glycemia. Diabetes 65, 34–43 (2016).

23. Kim, K. S., Seeley, R. J. & Sandoval, D. A. Signalling from the periphery to the brain that regulates energy homeostasis. Nature Reviews Neuroscience 19, 185–196 (2018).

24. Williams, E. K. K. et al. Sensory Neurons that Detect Stretch and Nutrients in the Digestive System. Cell 166, 209–221 (2016).

25. Bai, L. et al. Genetic Identification of Vagal Sensory Neurons That Control Feeding. Cell 179, 1129–1143.e23 (2019).

26. Holt, M. K. et al. Synaptic Inputs to the Mouse Dorsal Vagal Complex and Its Resident Preproglucagon Neurons. J. Neurosci. 39, 9767–9781 (2019).

27. Kreisler, A. D., Davis, E. A. & Rinaman, L. Differential activation of chemically identified neurons in the caudal nucleus of the solitary tract in non-entrained rats after intake of satiating vs. non-satiating meals. Physiol. Behav. 136, 47–54 (2014).

28. Nguyen, K. P., O’Neal, T. J., Bolonduro, O. A., White, E. & Kravitz, A. V. Feeding Experimentation Device (FED): A flexible open-source device for measuring feeding behavior. J. Neurosci. Methods 267, 108–114 (2016).

29. Kreisler, A. D. & Rinaman, L. Hindbrain glucagon-like peptide-1 neurons track intake volume and contribute to injection stress-induced hypophagia in meal-entrained rats. Am. J. Physiol. - Regul. Integr. Comp. Physiol. 310, R906–R916 (2016).

30. Ishii, Y. et al. Differential effects of the selective orexin-1 receptor antagonist SB-334867 and lithium chloride on the behavioural satiety sequence in rats. Physiol. Behav. 81, 129–140 (2004).

31. Wright, F. L. & Rodgers, R. J. Behavioural profile of exendin-4/naltrexone dose combinations in male rats during tests of palatable food consumption. Psychopharmacology (Berl). 231, 3729–3744 (2014).

32. Hisadome, K., Reimann, F., Gribble, F. M. & Trapp, S. Leptin directly depolarizes preproglucagon neurons in the nucleus tractus solitarius: Electrical properties of glucagon-like peptide 1 neurons. Diabetes 59, 1890–1898 (2010).

33. Vrang, N., Phifer, C. B., Corkern, M. M. & Berthoud, H.-R. Gastric distension induces c-Fos in medullary GLP-1/2-containing neurons. Am. J. Physiol. Integr. Comp. Physiol. 285, R470–R478 (2003).

34. Kanoski, S. E., Hayes, M. R. & Skibicka, K. P. GLP-1 and weight loss: unraveling the diverse neural circuitry. Am. J. Physiol. - Regul. Integr. Comp. Physiol. 310, R885–R895 (2016).

35. Yamaguchi, E., Yasoshima, Y. & Shimura, T. Systemic administration of anorexic gut peptide hormones impairs hedonic-driven sucrose consumption in mice. Physiol. Behav. 171, 158–164 (2017).

36. Hayes, M. R. & Schmidt, H. D. GLP-influences food and drug reward. Current Opinion in Behavioral Sciences 9, 66–70 (2016).

37. Card, J. P. et al. GLP-1 neurons form a local synaptic circuit within the rodent nucleus of the solitary tract. J. Comp. Neurol. 526, 2149–2164 (2018).

38. Anesten, F. et al. Preproglucagon neurons in the hindbrain have IL-6 receptor-α and show Ca2+ influx in response to IL-6. Am. J. Physiol. - Regul. Integr. Comp. Physiol. 311, R115–R123 (2016).

39. Holt, M. K., Llewellyn-Smith, I. J., Reimann, F., Gribble, F. M. & Trapp, S. Serotonergic modulation of the activity of GLP-1 producing neurons in the nucleus of the solitary tract in mouse. Mol. Metab. 6, 909–921 (2017).

40. Iwasaki, Y. et al. Peripheral oxytocin activates vagal afferent neurons to suppress feeding in normal and leptin-resistant mice: a route for ameliorating hyperphagia and obesity. Am. J. Physiol. - Regul. Integr. Comp. Physiol. 308, R360–R369 (2015).

41. Iwasaki, Y. et al. Relay of peripheral oxytocin to central oxytocin neurons via vagal afferents for regulating feeding. Biochem. Biophys. Res. Commun. 519, 553–558 (2019).

42. Kawatani, M., Yamada, Y. & Kawatani, M. Glucagon-like peptide-1 (GLP-1) action in the mouse area postrema neurons. Peptides 107, 68–74 (2018).

43. Yamamoto, H. et al. Glucagon-like peptide-1-responsive catecholamine neurons in the area postrema link peripheral glucagon-like peptide-1 with central autonomic control sites. J. Neurosci. 23, 2939–2946 (2003).

44. Hisadome, K., Reimann, F., Gribble, F. M. & Trapp, S. CCK stimulation of GLP-1 neurons involves α 1-adrenoceptor-mediated increase in glutamatergic synaptic inputs. Diabetes 60, 2701–2709 (2011).

45. Adams, J. M. et al. Liraglutide modulates appetite and body weight through glucagon-like peptide 1 receptor-expressing glutamatergic neurons. Diabetes 67, 1538–1548 (2018).

46. Fortin, S. M. et al. GABA neurons in the nucleus tractus solitarius express GLP-1 receptors and mediate anorectic effects of liraglutide in rats. Sci. Transl. Med. 12, eaay8071 (2020).

47. Llewellyn-Smith, I. J., Reimann, F., Gribble, F. M. & Trapp, S. Preproglucagon neurons project widely to autonomic control areas in the mouse brain. Neuroscience 180, 111–121 (2011).

48. Reimann, F. et al. Glucose Sensing in L Cells: A Primary Cell Study. Cell Metab. 8, 532–539 (2008).

49. Holt, M. K. et al. PPG neurons in the nucleus of the solitary tract modulate heart rate but do not mediate GLP-1 receptor agonist-induced tachycardia in mice. Mol. Metab. 101024 (2020). doi:10.1016/j.molmet.2020.101024

50. Parker, H. E. et al. Predominant role of active versus facilitative glucose transport for glucagon-like peptide-1 secretion. Diabetologia 55, 2445–2455 (2012).

51. Richards, P. et al. Identification and characterization of GLP-1 receptor-expressing cells using a new transgenic mouse model. Diabetes 63, 1224–33 (2014).

52. Han, W. et al. A Neural Circuit for Gut-Induced Reward. Cell 175, 665–678 (2018).

53. Tan, Y. et al. Oxytocin receptors are expressed by glutamatergic prefrontal cortical neurons that selectively modulate social recognition. J. Neurosci. 39, 3249–3263 (2019).

54. Rakipovski, G. et al. The GLP-1 Analogs Liraglutide and Semaglutide Reduce Atherosclerosis in ApoE −/− and LDLr −/− Mice by a Mechanism That Includes Inflammatory Pathways. JACC Basic to Transl. Sci. 3, 844–857 (2018).

55. de Lartigue, G., Ronveaux, C. C. & Raybould, H. E. Deletion of leptin signaling in vagal afferent neurons results in hyperphagia and obesity. Mol. Metab. 3, 595–607 (2014).

56. Friard, O. & Gamba, M. BORIS: a free, versatile open-source event-logging software for video/audio coding and live observations. Methods Ecol. Evol. 7, 1325–1330 (2016).

57. Halford, J. C., Wanninayake, S. C. & Blundell, J. E. Behavioral satiety sequence (BSS) for the diagnosis of drug action on food intake. Pharmacol. Biochem. Behav. 61, 159–68 (1998).

58. Rodgers, R. J., Holch, P. & Tallett, A. J. Behavioural satiety sequence (BSS): separating wheat from chaff in the behavioural pharmacology of appetite. Pharmacol. Biochem. Behav. 97, 3–14 (2010).

59. Mina, A. I. et al. CalR: A Web-Based Analysis Tool for Indirect Calorimetry Experiments. Cell Metab. 28, 656–666.e1 (2018).

60. de Kloet, A. D. et al. Reporter mouse strain provides a novel look at angiotensin type-2 receptor distribution in the central nervous system. Brain Struct. Funct. 221, 891–912 (2016).

61. Schindelin, J. et al. Fiji: An open-source platform for biological-image analysis. Nature Methods 9, 676–682 (2012).

62. Wu, Z., Autry, A. E., Bergan, J. F., Watabe-Uchida, M. & Dulac, C. G. Galanin neurons in the medial preoptic area govern parental behaviour. Nature 509, 325–30 (2014).

63. Krashes, M. J. et al. Rapid, reversible activation of AgRP neurons drives feeding behavior in mice. J. Clin. Invest. 121, 1424–8 (2011).

64. Watabe-Uchida, M., Zhu, L., Ogawa, S. K., Vamanrao, A. & Uchida, N. Whole-Brain Mapping of Direct Inputs to Midbrain Dopamine Neurons. Neuron 74, 858–873 (2012).

65. Wickersham, I. R. et al. Monosynaptic Restriction of Transsynaptic Tracing from Single, Genetically Targeted Neurons. Neuron 53, 639–647 (2007).

