## Supplemental figures for "Central and peripheral GLP-1 systems independently and additively suppress eating"

**Supplemental Information - Figures and Legends**

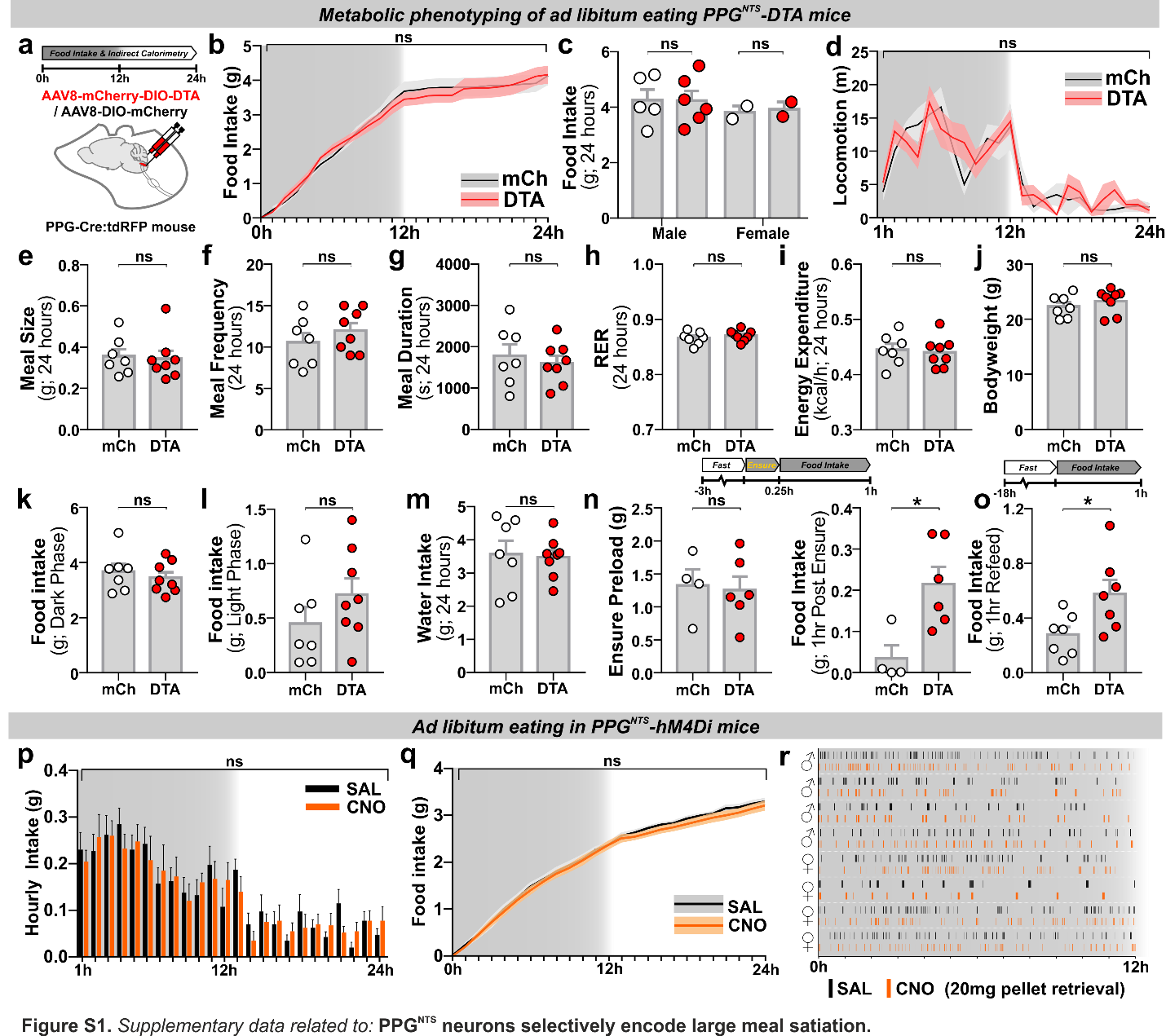

**Figure S1.** *Supplementary data related to:* **PPG^NTS^ neurons selectively encode large meal satiation.**

(a) Experimental model and paradigm for metabolic phenotyping of PPG^NTS^-DTA mice (DTA, n=8) or mCherry-transduced controls (mCh, n=7).

(b) Cumulative hourly food intake over 1 day, 2-way mixed-model ANOVA: Virus *F*_(1,13)_=0.015, *p*=0.904.

(c) Daily food intake by sex, 2-way mixed-model ANOVA: Virus *F*_(1,11)_=0.012, *p*=0.914; Sex *F*_(1,11)_=0.683, *p*=0.426.

(d) Directed ambulatory locomotion (excluding fine movements) over 1 day, 2-way mixed-model ANOVA: Virus *F*_(1,11)_=0.493, *p*=0.497.

(e-i) Meal pattern and metabolic parameters over 1 day, unpaired 2-tailed t-test or Mann-Whitney U test: e) *U*=25, *p*=0.779; f) *t*_(13)_=0.997, *p*=0.337; g) *t*_(13)_=0.565, *p*=0.582; h) *t*_(13)_=0.797, *p*=0.440; i) *t*_(13)_=0.323, *p*=0.752.

(j) Mean bodyweight over the 24h test period, unpaired 2-tailed t-test: *t*_(13)_=0.883, *p*=0.393.

(k-l) Food intake during dark and light phases, unpaired 2-tailed t-test: k) *t*_(13)_=0.668, *p*=0.516; l) *t*_(13)_=1.251, *p*=0.233.

(m) 24h water intake, unpaired 2-tailed t-test: *t*_(13)_=0.205, *p*=0.841.

(n) Ensure liquid diet preload intake, unpaired 2-tailed t-test: *t*_(8)_=0.219, *p*=0.832; and post-Ensure chow intake, Mann-Whitney U test: *U*=2, *p*=0.038.

(o) Post-fast refeed intake, unpaired 2-tailed t-test: *t*_(12)_=2.501, *p*=0.028.

(p-q) Hourly and cumulative intakes over 1 day from *ad libitum* eating PPG^NTS^-hM4Di mice (n=8), 2-way within-subjects ANOVA: p) Drug *F*_(1,7)_=0.241, *p*=0.639; q) Drug *F*_(1,7)_=0.411, *p*=0.542.

(r) Raster plot of chow pellet retrievals during the dark phase. Plots from the same mouse after saline and CNO injections presented adjacently.

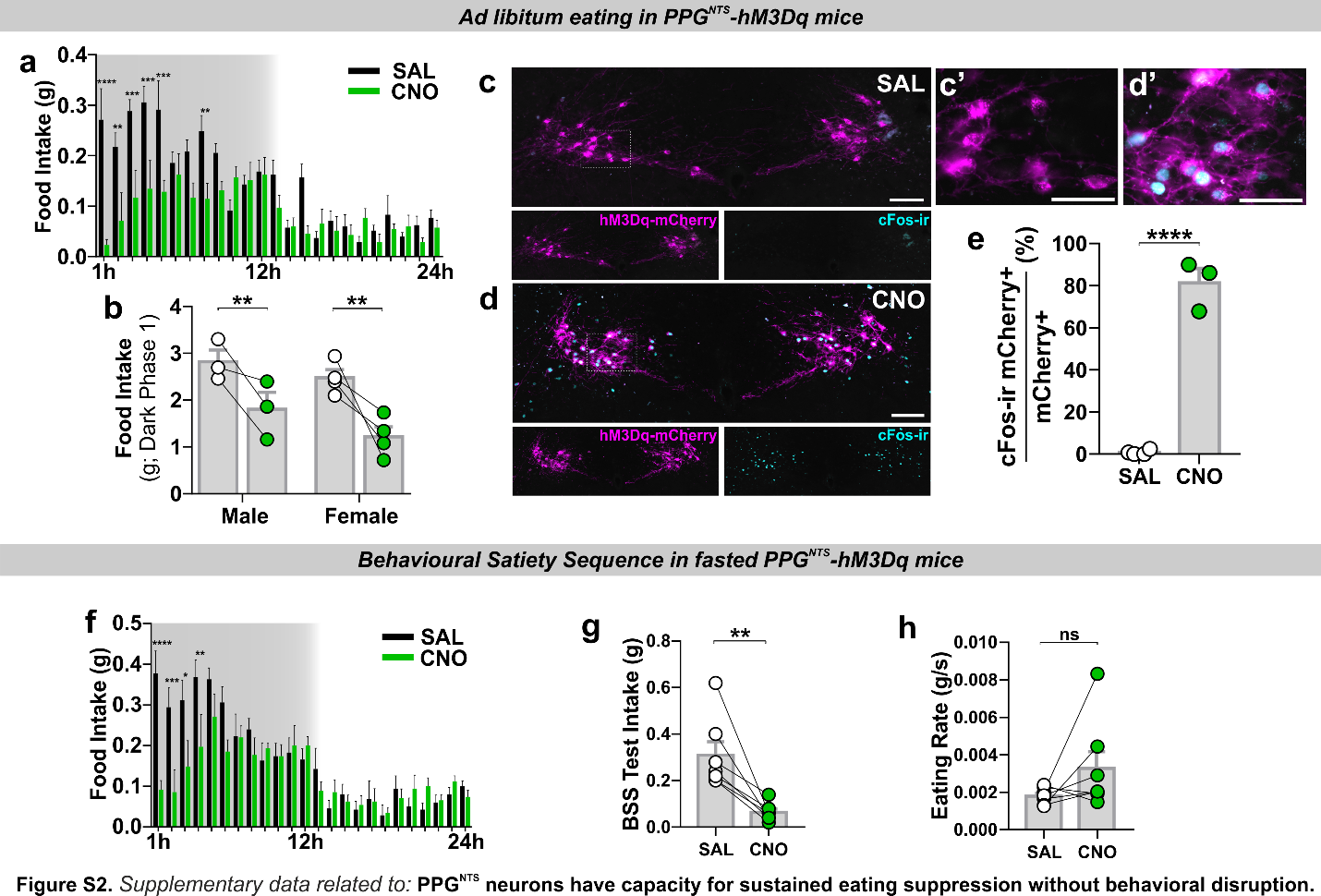

**Figure S2.** *Supplementary data related to:* **PPG^NTS^ neurons have capacity for sustained eating suppression without behavioural disruption.**

(a) Non-cumulative hourly food intake over the circadian cycle from *ad libitum* eating PPG^NTS^-hM3Dq mice (n=7), 2-way within-subjects ANOVA: Drug x Time *F*_(23,138)_=4.599, *p*<0.0001.

(b) Dark phase food intake by sex, 2-way mixed-model ANOVA: Drug *F*_(1,5)_=19.97, *p*=0.0066; Sex *F*_(1,5)_=3.854, *p*=0.107.

(c-e) Photomicrographs of co-localized cFos immunoreactivity and PPG^NTS^-hM3Dq-mCherry in NTS of mice perfused 3 hours after saline (c) or CNO (d) injection, and proportion of mCherry-expressing neurons co-localized with cFos-ir (e), unpaired 1-tailed t-test: *t*_(5)_=13.94, *p*<0.0001.

(f) Non-cumulative hourly food intake over 1 day from 18h fasted PPG^NTS^-hM3Dq mice (n=7), 2-way within-subjects ANOVA: Drug x Time *F*_(23,138)_=3.745, *p*<0.0001. The behavioural satiety sequence (BSS) was analysed during the first 40 minutes of the dark phase.

(g-h) Food intake and eating rate over 40 minute BSS test, paired 2-tailed t-test and Wilcoxon matched pairs test: g) *t*_(6)_=4.088, *p*=0.0064; h) W=18, *p*=0.156.

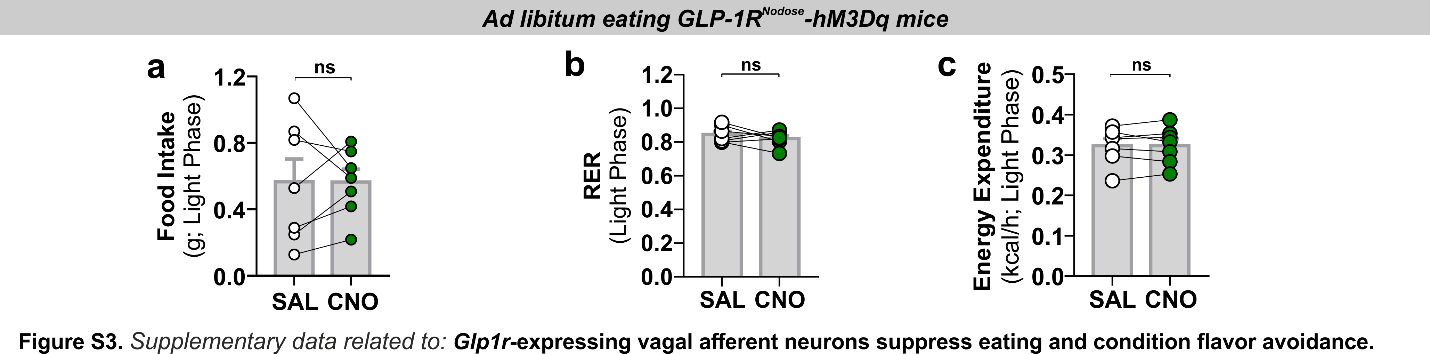

**Figure S3.** *Supplementary data related to:* ***Glp1r*-expressing vagal afferent neurons suppress eating and condition flavour avoidance.**

(a-c) Light phase food intake and metabolic parameters from *ad libitum* eating GLP-1R^Nodose^-hM3Dq mice, paired 2-tailed t-test: a) *t*_(6)_=0.0141, *p*=0.989; b) *t*_(6)_=0.952, *p*=0.378; c) *t*_(6)_=0.0406, *p*=0.969.

**Figure S4.** *Supplementary data related to:* ***Oxtr* rather than *Glp1r* VANs are the major vagal input to PPG^NTS^ neurons*.***
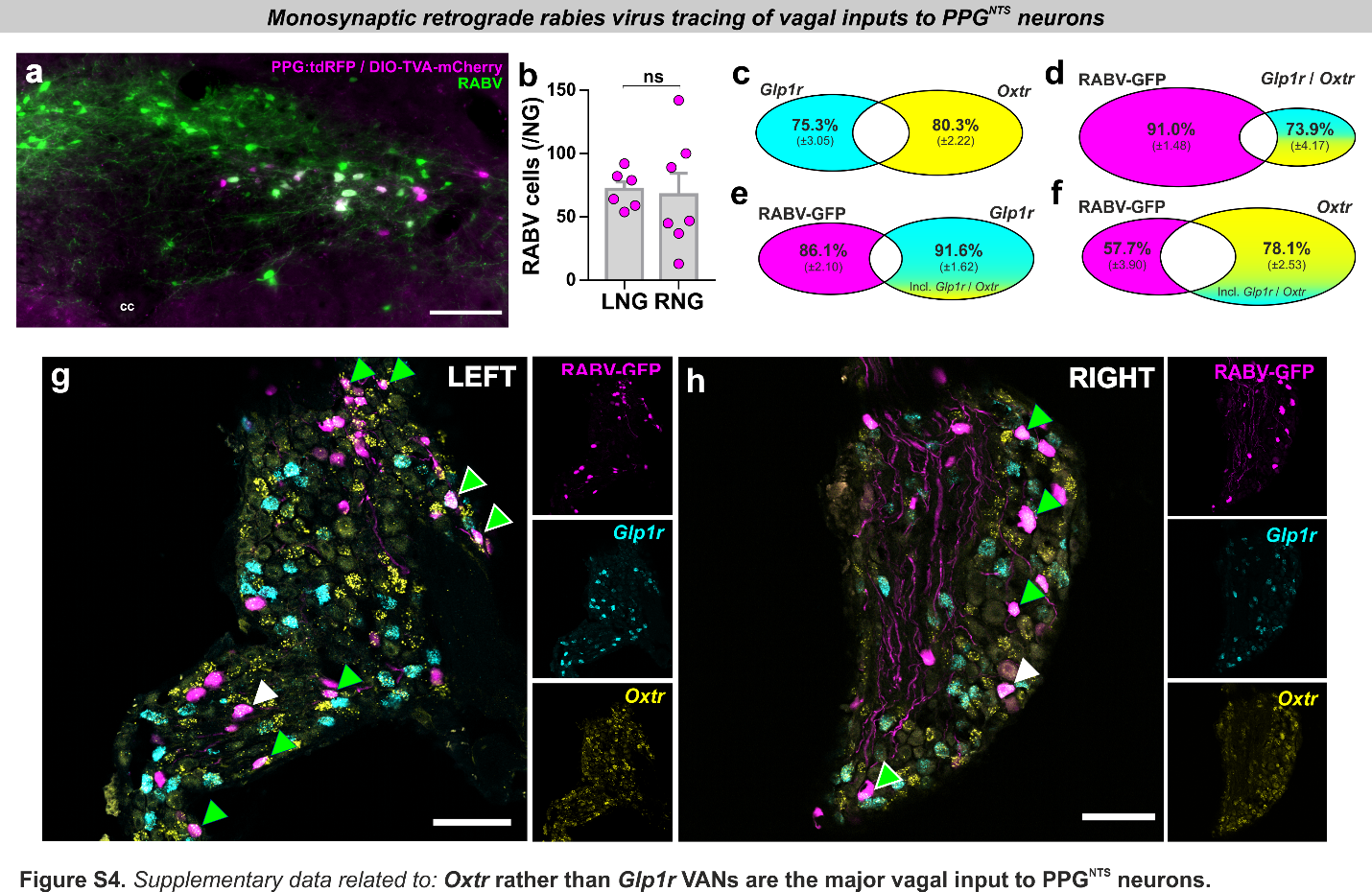

(a) Representative photomicrograph of coronal NTS section from PPG-Cre:tdRFP mouse transduced with DIO-TVA-mCherry + DIO-RabiesG, and subsequently with rabies virus-ΔG-GFP (RABV). Bilateral NTS injection of TVA+RabiesG and counterbalanced unilateral injection of RABV (4 mice / side) resulted in 40.5% (±5.5) of all PPG^NTS^ neurons being successfully transduced ‘starter’ neurons, identified by co-localization of mCherry (and/or tdRFP) and GFP. Despite unilateral RABV injection, starter neurons were observed in left and right NTS in all mice, indicating significant viral spread and bilateral transduction.

(b) Total RABV+ cells in left and right nodose ganglia (LNG / RNG), unpaired 2-tailed t-test: *t*_(11)_=0.214, *p*=0.834.

(c-d) Quantification of *Glp1r* and *Oxtr* co-localization in nodose ganglia (c), and proportions of dual-expressing *Glp1r* / *Oxtr* cells co-localized with RABV (d). This dual population comprises 24.7% of all *Glp1r* cells and 19.7% of all *Oxtr* cells. 9% of RABV+ vagal inputs to PPG^NTS^ neurons express both *Glp1r* and *Oxtr*, and 26.1% of dual-expressing *Glp1r* / *Oxtr* cells are RABV+ vagal inputs to PPG^NTS^ neurons.

(e) Quantification of RABV and *Glp1r* co-localization in NG as proportions of all RABV+ cells and all *Glp1r*+ cells, including those *Glp1r* cells that also express *Oxtr*.

(f) Quantification of RABV and *Oxtr* co-localization in NG as proportions of all RABV+ cells and all *Oxtr*+ cells, including those *Oxtr* cells that also express *Glp1r*.

(g-h) Photomicrographs of left and right nodose ganglion sections showing rabies virus GFP expression (RABV) and *Glp1r* and *Oxtr* FISH. RABV+*Glp1r* co-localization shown by white arrows, RABV+*Oxtr* by green arrows and RABV+*Glp1r*+*Oxtr* by white-edged green arrow. Scale=100μm.

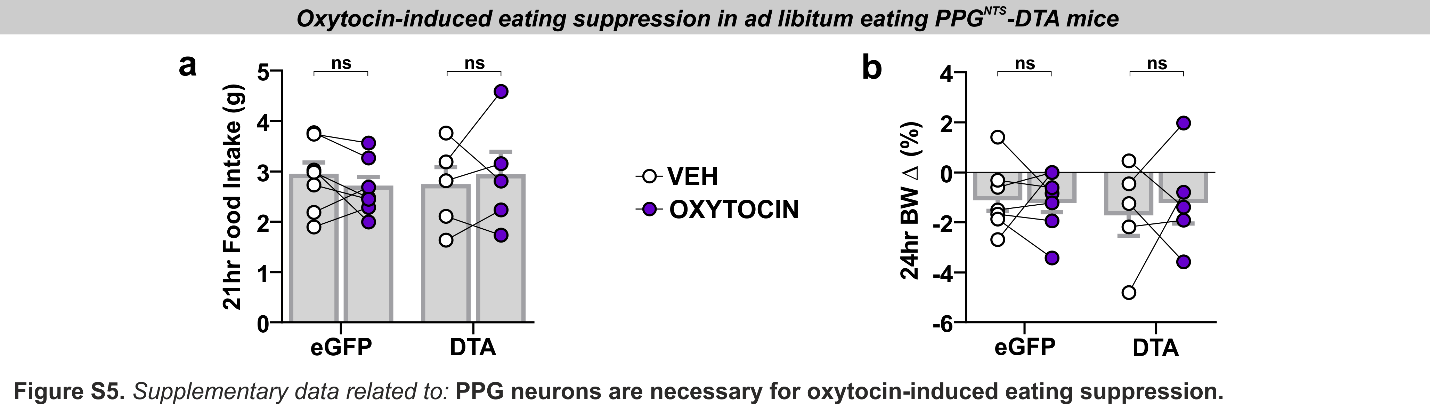

**Figure S5.** *Supplementary data related to:* **PPG^NTS^ neurons are necessary for oxytocin-induced eating suppression.**

(a-b) Food intake and bodyweight change over 1 day in eGFP and DTA mice administered oxytocin (0.4 mg/kg, i.p.), 2-way mixed-model ANOVA: a) Drug *F*_(1,10)_=0.00474, *p*=0.947; b) Drug *F*_(1,10)_=0.0989, *p*=0.760.

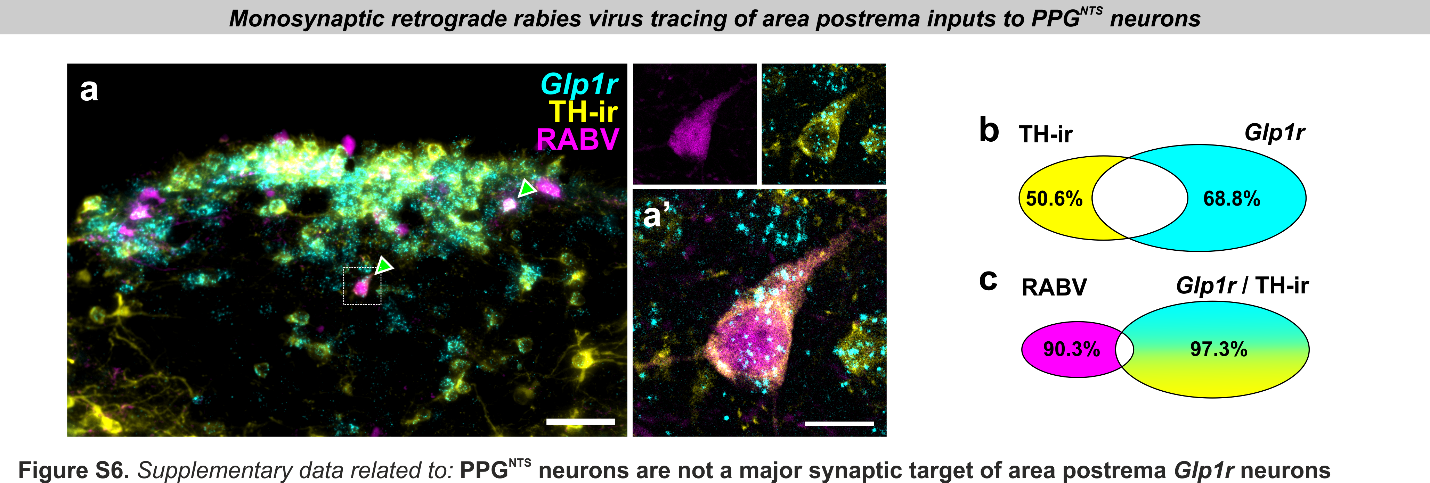

**Figure S6.** *Supplementary data related to:* **PPG^NTS^ are not a major synaptic target of area postrema *Glp1r* neurons.**

(a) Photomicrographs of coronal NTS section showing RABV expression, *Glp1r* FISH and TH-ir. RABV+*Glp1r*+TH-ir co-localization shown by white-edged green arrows. Scale=100μm (inset 20μm).

(b-c) Quantification of *Glp1r* and TH-ir co-localization in area postrema (b), and proportions of dual *Glp1r* / TH-ir cells co-localized with RABV (c). This dual population comprises 49.4% of all TH-ir cells and 31.2% of all *Glp1r* cells. 9.7% of RABV+ AP inputs to PPG^NTS^ neurons express *Glp1r* and are TH-ir, and 2.7% of dual *Glp1r* / TH-ir cells are RABV+ AP inputs to PPG^NTS^ neurons.

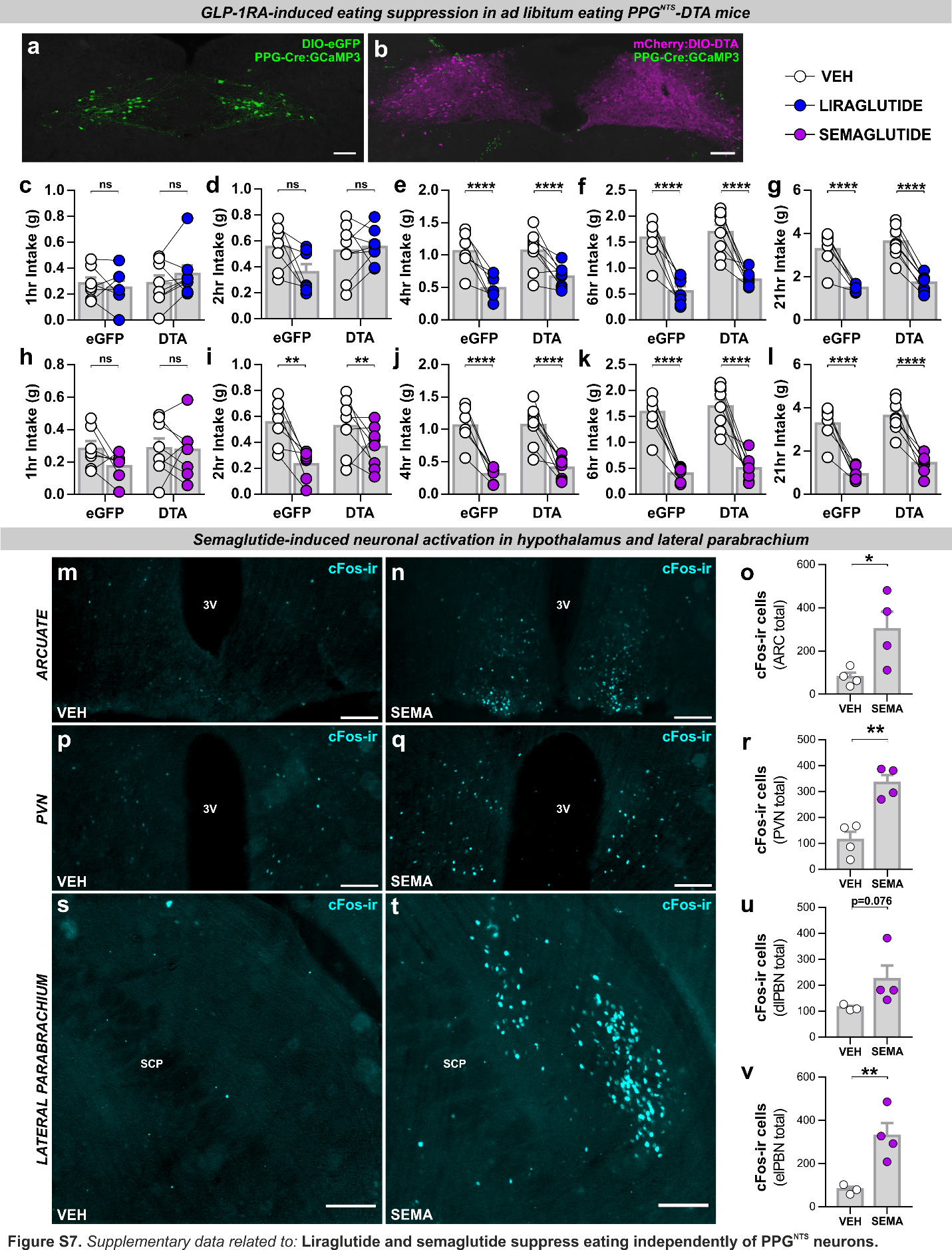

**Figure S7.** *Supplementary data related to:* **Liraglutide and semaglutide suppress eating independently of PPG^NTS^ neurons.**

(a-b) Photomicrographs of coronal NTS sections from PPG-Cre:GCaMP3 mice injected with eGFP control virus (a) or DTA virus (b). Note the complete absence of green (GCaMP3-expressing) PPG^NTS^ neurons in DTA-ablated tissue, and extent of viral spread as demonstrated by Cre-independent expression of mCherry.

(c-g) Cumulative food intake by virus at 1,2,4,6 and 21hr in eGFP and DTA mice administered liraglutide (200 μg/kg, s.c.), 2-way mixed-model ANOVA: c) Drug *F*_(1,13)_=0.246, *p*=0.628; d) Drug *F*_(1,13)_=2.108, *p*=0.170; e) Drug *F*_(1,13)_=37.44, *p*<0.0001, Virus *F*_(1,13)_=0.836, *p*=0.377; f) Drug *F*_(1,13)_=75.09, *p*<0.0001, Virus *F*_(1,13)_=1.877, *p*=0.194; g) Drug *F*_(1,13)_=154.9, *p*<0.0001, Virus *F*_(1,13)_=1.272, *p*=0.280.

(h-l) Cumulative food intake by virus at 1,2,4,6 and 21hr in eGFP and DTA mice administered semaglutide (60 μg/kg, s.c.), 2-way mixed-model ANOVA: h) Drug *F*_(1,13)_=1.965, *p*=0.184; i) Drug *F*_(1,13)_=17.1, *p*=0.0012; Virus *F*_(1,13)_=0.630, *p*=0.442; j) Drug *F*_(1,13)_=82.49, *p*<0.0001, Virus *F*_(1,13)_=0.332, *p*=0.574; k) Drug *F*_(1,13)_=98.21, *p*<0.0001, Virus *F*_(1,13)_=0.840, *p*=0.376; l) Drug *F*_(1,13)_=126.1, *p*<0.0001, Virus *F*_(1,13)_=3.42, *p*=0.0873.

(m-o) Photomicrographs of cFos immunoreactivity (cFos-ir) in arcuate nucleus of the hypothalamus (ARC) 4 hours after vehicle (VEH; n=4) or semaglutide (SEMA, 60 μg/kg, s.c.; n=4) administration, and total cFos count, unpaired 1-tailed t-test: o) *t*_(6)_=2.614, *p*=0.020. Scale=100μm.

(p-r) Photomicrographs of cFos-ir in paraventricular nucleus of the hypothalamus (PVN) 4 hours after vehicle or semaglutide administration (both n=4), and total cFos count, unpaired 1-tailed t-test: r) *t*_(6)_=5.109, *p*=0.0011. Scale=100μm.

(s-v) Photomicrographs of cFos-ir in dorsal lateral and external lateral subdivisions of the parabrachial nucleus (dlPBN / elPBN) 4 hours after vehicle or semaglutide administration (n=3/4), and total cFos count, unpaired 1-tailed t-tests: u) *t*_(5)_=1.693, *p*=0.0756; v) *t*_(5)_=3.57, *p*=0.0080. Semaglutide did not increase cFos-ir in the medial PBN (data not shown), *t*_(5)_=0.435, *p*=0.341. Scale=100μm.
